# A sustained type I IFN-neutrophil-IL-18 axis drives pathology during mucosal viral infection

**DOI:** 10.1101/2020.12.20.423690

**Authors:** Tania J. Lebratti, Ying Shiang Lim, Adjoa Cofie, Prabhakar S. Andey, Xiaoping Jiang, Jason M. Scott, Maria Rita Fabbrizi, Ayse N. Ozanturk, Christine T.N. Pham, Regina A. Clemens, Maxim Artyomov, Mary C. Dinauer, Haina Shin

## Abstract

Neutrophil responses against pathogens must be balanced between protection and immunopathology. Factors that determine these outcomes are not well-understood. In a mouse model of genital herpes simplex virus-2 (HSV-2) infection, which results in severe genital inflammation, antibody-mediated neutrophil depletion reduced disease. Comparative single cell RNA-sequencing analysis of vaginal cells against a model of genital HSV-1 infection, which results in mild inflammation, demonstrated sustained expression of interferon-stimulated genes (ISGs) only after HSV-2 infection primarily within the neutrophil population. Both therapeutic blockade of IFN*α/β* receptor 1 (IFNAR1) and genetic deletion of IFNAR1 in neutrophils concomitantly decreased HSV-2 genital disease severity and vaginal IL-18 levels. Therapeutic neutralization of IL-18 also diminished genital inflammation, indicating an important role for this cytokine in promoting neutrophil-dependent immunopathology. Our study reveals that sustained type I IFN signaling is a driver of pathogenic neutrophil responses, and identifies IL-18 as a novel component of disease during genital HSV-2 infection.

## INTRODUCTION

Neutrophils are a critical component of the innate immune system. In humans, they are the most abundant leukocyte in circulation and are often amongst the first wave of immune cells responding to pathogen invasion. In the context of bacterial or fungal infection, including those that are sexually transmitted, neutrophils are largely protective and can help eliminate pathogens through a variety of effector functions, including phagocytosis, production of reactive oxygen species (ROS), NET and protease release, and cytokine and chemokine secretion (Mayadas, Cullere, & Lowell, 2014; Pham, 2006; Tecchio, Micheletti, & Cassatella, 2014). In contrast, the role of neutrophils during viral infection is less clear (Galani & Andreakos, 2015). While neutrophils have been reported to neutralize several viruses and display protective qualities in vivo (Akk, Springer, & Pham, 2016; Craig N. Jenne et al., 2013; Saitoh et al., 2012; Tate et al., 2009; Tate et al., 2011) they have also been associated with tissue damage, loss of viral control, and increased mortality (Bai et al., 2010; Brandes, Klauschen, Kuchen, & Germain, 2013; Kulkarni et al., 2019; Narasaraju et al., 2011; Vidy et al., 2016).

Type I IFNs are potent regulators of neutrophil activity in a multitude of contexts. Type I IFN can enhance recruitment of neutrophils to sites of infection, regulate neutrophil function and drive immunopathology after infection by different classes pathogens, including *Plasmodium spp*., *Candida spp*. and *Pseudomonas spp*. (Majer et al., 2012; Pylaeva et al., 2019; Rocha et al., 2015). However, type I IFNs can also inhibit neutrophil recruitment to the ganglia by suppressing chemokine expression after HSV infection (Stock, Smith, & Carbone, 2014), suggesting that the interplay of IFNs and neutrophil activity may be dependent on tissue type and the pathogen. The relationship between neutrophil-intrinsic type I IFN signaling and infection outcomes is less clear. Type I IFNs can promote expression of interferon stimulated genes (ISGs) and pro-inflammatory cytokines in neutrophils, suggesting a potential role in driving immunopathology (Galani et al., 2017). During Leishmania infection, however, IFNAR signaling appears to suppress neutrophil-dependent killing of parasites (Xin et al., 2010), which emphasizes the complexity of IFN-mediated neutrophil responses.

Genital herpes is a chronic, sexually transmitted infection that affects over 400 million people worldwide (World Health Organization, 2007) and can be caused by two members of the alphaherpesvirus family, herpes simplex virus-2 (HSV-2) or the related HSV-1. Genital herpes is characterized by recurrent episodes of inflammation and ulceration and the factors that drive disease are unclear. In humans, ulcer formation is associated with suboptimal viral control and spread during episodes of reactivation (Roychoudhury et al., 2020; J. T. Schiffer & Corey, 2013; Joshua T Schiffer et al., 2013), while in mouse models, severity of disease often correlates with susceptibility to infection and the level of viral replication in the genital mucosa (Gopinath et al., 2018). Neutrophil infiltration into sites of active HSV-2 ulcers have also been reported in humans (Boddingius, Dijkman, Hendriksen, Schift, & Stolz, 1987), but whether these cells are helpful or harmful during HSV infection is unknown. While neutrophils have been associated with tissue damage after multiple routes of HSV-1 infection (Divito & Hendricks, 2008; Khoury-Hanold et al., 2016; Rao & Suvas, 2018; Thomas, Gangappa, Kanangat, & Rouse, 1997), a protective role for neutrophils after genital HSV-2 infection has also been reported (Milligan, 1999; Milligan, Bourne, & Dudley, 2001), although use of non-specific depletion antibodies have muddled the respective contribution of neutrophils and other innate immune cells such as monocytes, which are known to be antiviral (Iijima, Mattei, & Iwasaki, 2011). Furthermore, increased neutrophil recruitment to the HSV-2 infected vaginal epithelial barrier resulted in greater epithelial cell death, suggesting that neutrophil responses may indeed be pathogenic (Krzyzowska et al., 2014). However, the factors that distinguish pathogenic vs non-pathogenic neutrophil responses during viral infection, including HSV-2 infection, remain ill-defined.

To address this, we evaluated the impact of neutrophils on genital disease severity using two models of HSV infection that result in low levels (HSV-1) or high levels of inflammation (HSV-2) (A. G. Lee et al., 2020). Between these two states, heightened expression of type I IFN during the resolution phase of acute infection and sustained expression of ISGs in neutrophils was detected after HSV-2 infection but not HSV-1. Therapeutic antibody-mediated blockade of IFNAR1 as well as neutrophil-specific deletion of IFNAR1 reduced both genital inflammation as well as vaginal IL-18 levels during the resolution phase of acute HSV-2 infection. Accordingly, therapeutic neutralization of IL-18 also ameliorated genital disease after HSV-2 infection. Together, our data demonstrates that sustained type I IFN signaling is a key determinant of pathogenic neutrophil responses during viral infection, and identifies neutrophil- and type I IFN-dependent IL-18 production as a novel driver of inflammation during genital HSV-2 infection.

## RESULTS

### Neutrophils are a component of severe genital inflammation after vaginal HSV-2 infection

To determine the role of neutrophils in our model of vaginal HSV-2 infection, wildtype (WT) female C57BL/6 mice were treated with Depo-Provera (depot medroxyprogestrone, DMPA) to hold mice at the diestrus phase of the estrus cycle and normalize susceptibility to infection (Kaushic, Ashkar, Reid, & Rosenthal, 2003). Neutrophils were depleted in DMPA-treated mice by intraperitoneal (i.p.) injection of an antibody against Ly6G, a neutrophil-specific marker, or an isotype control. One day later, mice were inoculated intravaginally with 5000 plaque forming units (PFU) of WT HSV-2 strain 186 syn+ (WT HSV-2). Neutrophils were effectively depleted from the vagina up to 5 d.p.i. (Figure 1A). In order to focus on genital inflammation, mice were monitored for 1 week after infection, as progression of disease within the second week of our infection model is largely indicative of viral dissemination into the central nervous system. In both cohorts, mild genital inflammation was apparent starting at 4 d.p.i. in a small fraction of mice (Figure 1B). Over time, progression of disease in the neutrophil-depleted mice was significantly slower compared to the controls. Remarkably, as late as 7 d.p.i., a proportion of the neutrophil-depleted group remained uninflamed, in contrast to the isotype control group in which all mice displayed signs of inflammation (Figure 1B). To confirm the disparity in scored disease severity, we examined the vagina and genital skin by histology. At 6 d.p.i., epithelial denuding and damage was apparent in the isotype control-treated mice (Figure 1C). In contrast, only a limited amount of epithelial destruction was observed in neutrophil-depleted mice, with less cellular infiltrates at sites of damage and in the lumen (Figure 1C). Furthermore, the epithelial layer proximal to areas of damage was morphologically distinct in isotype control-treated animals compared to neutrophil-depleted animals, suggesting diverse epithelial responses after infection in the presence or absence of neutrophils (Figure 1C). Similarly, destruction of the epidermis and separation of the epidermis from the dermis was widespread in the genital skin of isotype control-treated mice but not in neutrophil-depleted mice (Figure 1C). Unexpectedly, differences in genital inflammation and mucosal damage were largely independent of changes in viral control in the absence of neutrophils, as viral shedding into the vaginal lumen was similar between the two groups (Figure 1D). Indeed, disease severity was decreased in neutrophil-depleted mice despite a slight delay in the resolution of viral replication at 5 d.p.i. (Figure 1D).

**Figure 1.**
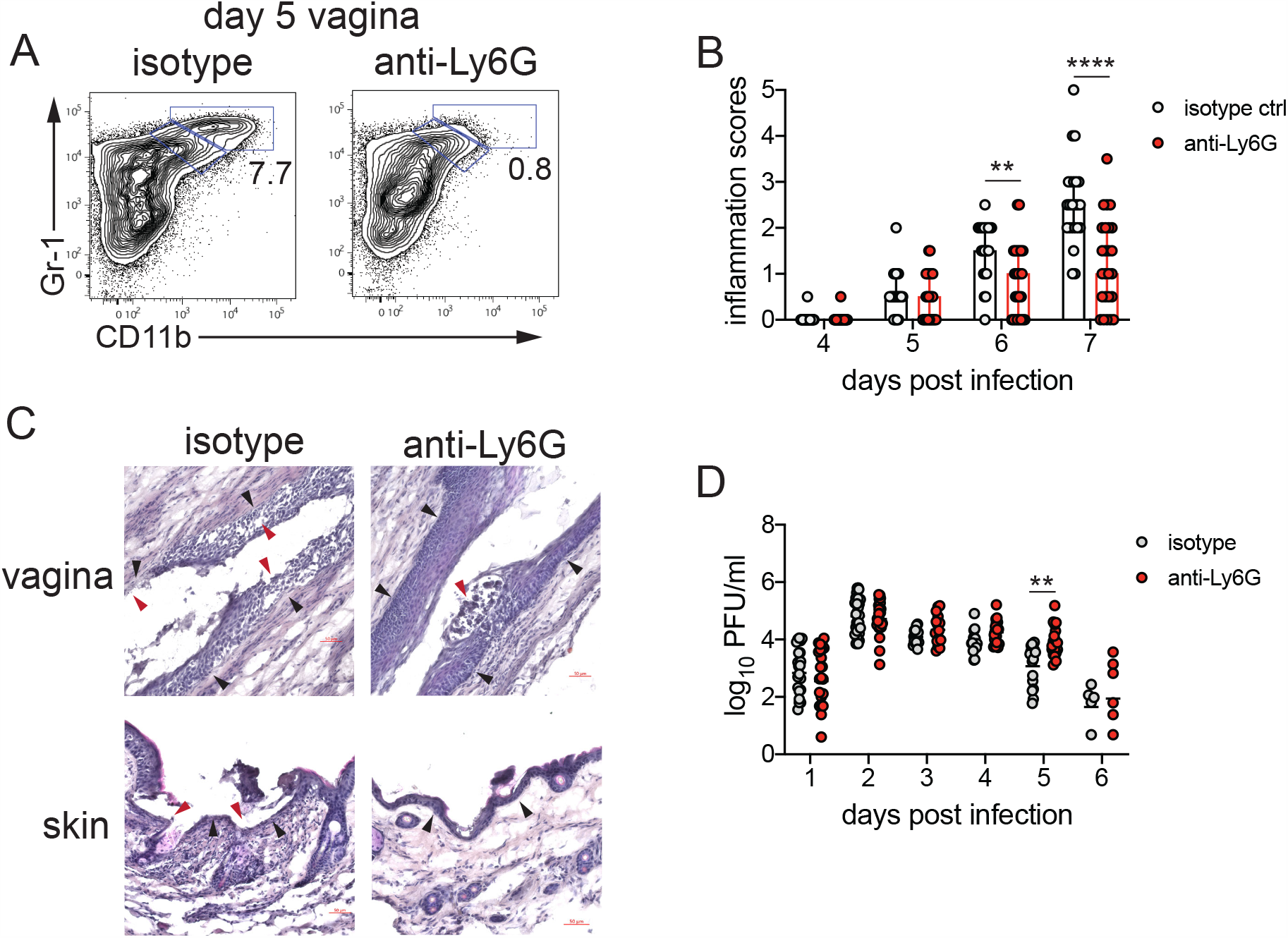
Neutrophil depletion reduces disease severity after HSV-2 vaginal infection. Female C57BL/6J mice were treated with DMPA and inoculated intravaginally (ivag) with 5000 PFU HSV-2. One day prior to HSV-2 inoculation, mice were injected intraperitoneally (i.p.) with 500μg of rat IgG2a isotype control or anti-Ly6G. **A**. Depletion was confirmed by flow cytometry in the vagina at 5 d.p.i.. Numbers in plots show the frequency of Gr-1+ CD11b+ neutrophils. **B**. Inflammation scores over the first 7.d.p.i. of mice treated with anti-Ly6G antibody (n=25) or isotype control (n=23) Mice showed no signs of disease prior to 4 d.p.i.. **C**. Histology of the vagina (top) or genital skin (bottom) at 6 d.p.i. from isotype control (left) or anti-Ly6G antibody-treated mice (right). Red arrows point to areas of epithelial denuding or damage, black arrows denote the basement membrane. **D**. Infectious virus as measured by plaque assay in vaginal washes collected daily (both groups day 1: n=22, day 2: n=28, day 3: n=15, day 4: n=16, day 5: n=19, day 6: n=6). Data in **B** and **D** are pooled from 4 independent experiments. Data in **C** is representative of 2 independent experiments. Bars in **B** show median with interquartile range. Horizontal bars in **C** show mean. Statistical analysis was performed by repeated measures two-way ANOVA with Geisser-Greenhouse correction and Bonferroni’s multiple comparisons test (**B**) or repeated measures two-way ANOVA with Bonferroni’s multiple comparison’s test (**D**). **p<0.01, ****p<0.001. Raw values for each biological replicate, epsilon values and specific p values are provided in Figure 1 - Source Data.

We next evaluated whether the decreased inflammation after neutrophil depletion was due to changes in the cellular response against HSV-2 infection. We examined the recruitment of Ly6C+ monocytes, NK cells, CD4 and CD8 T cells, all of which have been implicated in either the control of HSV or modulation of disease severity (A. J. Lee & Ashkar, 2012; Shin & Iwasaki, 2013; Truong, Smith, Sandgren, & Cunningham, 2019). To remove intravascular cells and to limit our analysis to cells within the vagina, tissues were thoroughly perfused prior to collection (Scott et al., 2018). Unexpectedly, there was no significant difference in the number of Ly6C+CD11b+ cells (Figure 1 - Supplement 1A), NK cells (Figure 1 - Supplement 1B), total CD4 (Figure 1 - Supplement 1C) or CD8 T cells (Figure 1 - Supplement 1D) that were recruited to the vagina over the first six days after infection regardless of whether neutrophils were present or not. Thus, our data demonstrate that neutrophils do not play a significant antiviral role in our model of vaginal HSV-2 infection, and rather promote genital inflammation with minimal impact viral burden and recruitment of other immune cells to the vagina.

### Neutrophil extracellular trap formation and oxidative burst are not major drivers of genital inflammation after HSV-2 infection

We next wanted to determine whether neutrophil-specific effector functions were promoting disease after HSV-2 infection. NETs have been associated with tissue damage in the context of both infectious (C. N. Jenne & Kubes, 2015) and non-infectious disease (Granger, Peyneau, Chollet-Martin, & de Chaisemartin, 2019). To test whether NETs play a role in genital disease after HSV-2 infection, we first examined the ability of neutrophils to form NETs when exposed to HSV-2. *In vitro* stimulation of neutrophils with HSV-2 resulted the enlargement of cell nuclei and the characteristic expulsion of DNA coated in citrullinated histones, which is a key characteristic of NETs (Figure 1 - Supplement 2A). The formation of NETs requires input from multiple pathways, including histone citrullination by enzymes such as PAD4, which leads to chromatin de-condensation and the eventual release of DNA (P. Li et al., 2010). To generate animals that were specifically lacking PAD4 in neutrophils, we bred PAD4^fl/fl^ x MRP8-Cre mice (PAD4 CKO). HSV-2 infection of these mice and their littermate controls demonstrated minimal impact on genital inflammation (Figure 1 - Supplement 2B) or viral replication (Figure 1 - Supplement 2C) in the genital mucosa. Thus, our data show PAD4 expression in neutrophils, and likely NET formation, are not the mechanisms by which these cells mediate disease after HSV-2 infection.

We next tested whether ROS production by neutrophils mediated inflammation after HSV-2 infection. While production of ROS in neutrophils supports antimicrobial activity against a variety of pathogens (Dinauer, 2019), excessive oxidative stress can be associated with tissue injury (Mittal, Siddiqui, Tran, Reddy, & Malik, 2014). We found that *in vitro* stimulation of neutrophils with HSV-2 led to an increase in ROS production compared to unstimulated cells (Figure 1 - Supplement 3A). To determine whether respiratory burst in neutrophils promoted genital inflammation after HSV-2 infection *in vivo*, we infected mice with germline deficiency in Ncf2 (Ncf2 KO), which encodes p67^*phox*^, a key component of the NADPH oxidase complex (Jacob et al., 2017). HSV-2 infection of Ncf2 KO and heterozygous controls resulted in similar progression of disease (Figure 1 - Supplement 3B) and did not alter viral titer (Figure 1 - Supplement 3C). To confirm that tested neutrophil effector functions, including ROS production, we infected mice in which the calcium-sensing molecules STIM1 and STIM2 were deleted from neutrophils (Clemens, Chong, Grimes, Hu, & Lowell, 2017). Stim1^fl/fl^ x Stim2^fl/fl^ x MRP8-Cre (STIM1/2 DKO) mice were infected with HSV-2 and monitored for disease. As expected, there was little difference in genital inflammation severity between the STIM1/2 DKO and Cre-controls (Figure 1- Supplement 3D) or viral titers (Figure 1 - Supplement 3E). Together, our data show that ROS production from neutrophils and other cell types play little role in driving genital inflammation after HSV-2 infection.

### A type I IFN signature distinguishes neutrophil responses after genital HSV-1 and HSV-2 infection

To identify the factors that drove pathogenic neutrophil responses after HSV-2 infection, we turned to a complementary model of HSV-1 genital infection that we had previously described (A. G. Lee et al., 2020). Inoculation with the same dose of HSV-1 and HSV-2 led to profound differences in genital inflammation (Figure 2A) despite comparable levels of mucosal viral shedding (A. G. Lee et al., 2020). Importantly, magnitude of the neutrophil response in the vagina was similar between HSV-1 and HSV-2 infected mice during the course of acute mucosal infection (Figure 2B), and neutrophils could be found infiltrating sites of both HSV-1 and HSV-2-infected epithelium, although it appeared that the interaction between neutrophil and virally infected epithelial cells after HSV-1 inoculation was not as extensive as after HSV-2 inoculation (Figure 2C). In contrast to HSV-2 infection, antibody-mediated depletion of neutrophils with anti-Ly6G antibody prior to inoculation with HSV-1 did not reduce the development of genital inflammation during the first 7 days after infection (Figure 2 - Supplement 1). Together, our data suggests that the regulation of the neutrophil response after HSV-1 or HSV-2 infection was distinct, leading to disparate inflammatory outcomes.

**Figure 2.**
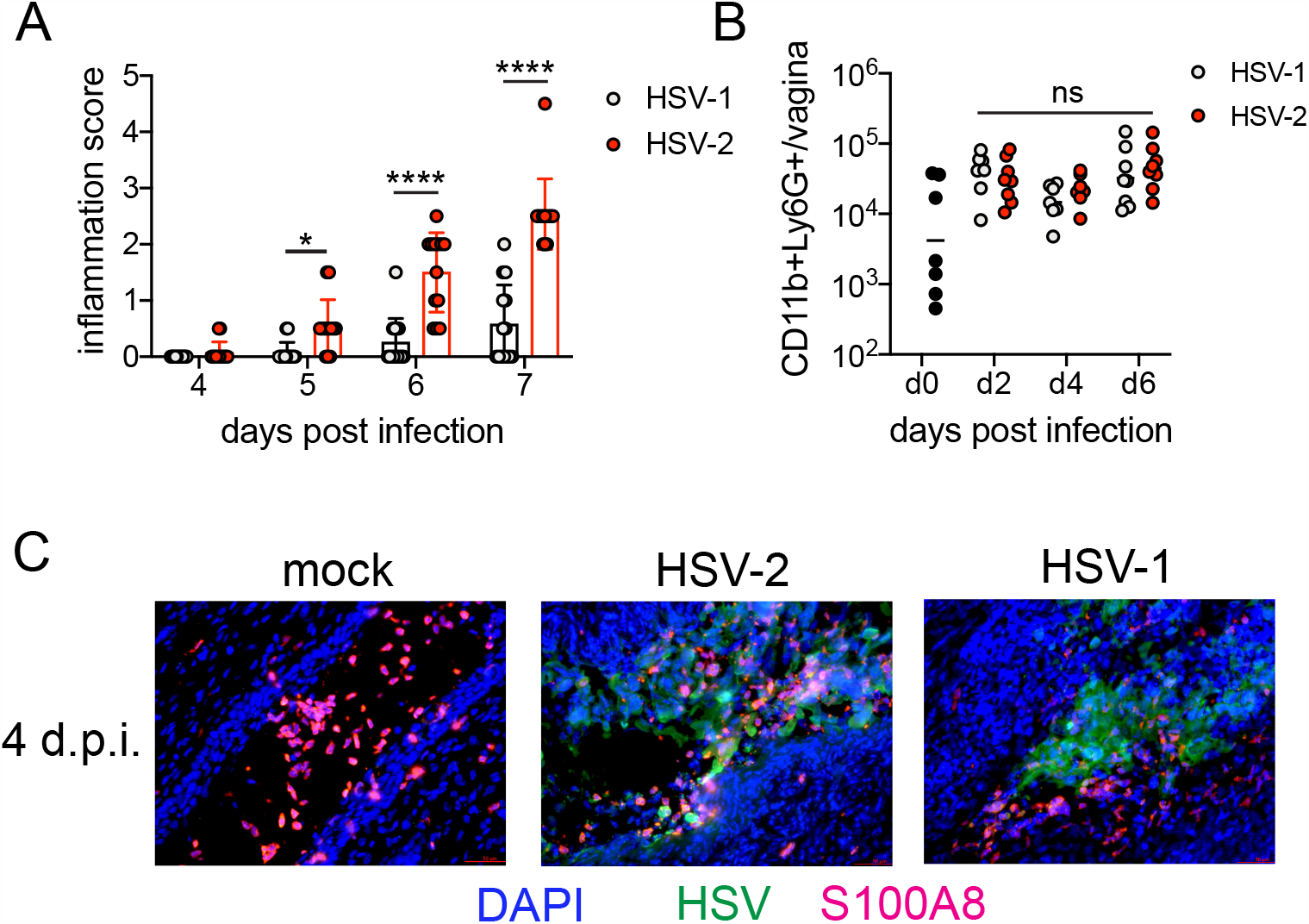
Magnitude of the neutrophil response during HSV-1 and HSV-2 is similar despite difference in disease outcome. Female C57BL/6J mice were treated with DMPA and inoculated ivag with 10^4^ PFU HSV-1 McKrae or HSV-2. **A**. Inflammation scores were monitored for 7 d.p.i.. For HSV-1: n=14, HSV-2: n=13. **B**. Neutrophils were counted by flow cytometry in vaginal tissues at the indicated days after HSV-1 or HSV-2 infection. For d0: n=7, day 2: n=8, day 4: n=7, day 6: n=8. **C**. Vaginas were harvested from PBS inoculated (mock), HSV-1 or HSV-2 inoculated mice at 4 d.p.i., and tissue sections were probed with antibodies against HSV proteins (green) or S100A8 (red). DAPI (blue) was used detect cell nuclei. Images are representative of six mice per group. Data are pooled from 3 (**A**) or 2 (**B, C**) independent experiments. Bars show median with interquartile range in **A** and mean in **B**. Statistical significance was measured by repeated measures two-way ANOVA with Geisser-Greenhouse correction and Bonferroni’s multiple comparisons test (**A**) or two-way ANOVA with Bonferroni’s multiple comparisons test (**B**). *p<0.05, ****p<0.001, ns = not significant. Raw values for each biological replicate, epsilon values and specific p values are provided in Figure 2 - Source Data.

To better better understand the differences between pathogenic neutrophil responses after HSV-2 infection and the non-pathogenic neutrophil responses after HSV-1, we performed single cell RNA-sequencing (scRNA-seq) on sorted live vaginal cells from a mock infected mouse or mice infected with HSV-1 or HSV-2 using the 10x Genomics platform (Zheng et al., 2017). Each sample was comprised of cells from a single animal in to better delineate potential subsets within cell populations, particularly neutrophils. Analysis across 21,633 cells in all samples revealed 17 unique clusters in the vagina during HSV infection after filtering, including myeloid cells, lymphocytes and epithelial cells (Figure 3A). Neutrophils were identified by expression of known cell markers such as *S100a8* and *Csf3r* (Figure 3B). In mock infected animals, the vaginal neutrophil population was dominated by cluster 0, and upon infection, at least two additional neutrophil subsets, cluster 2 and cluster 5, were clearly present (Figure 3C). While HSV-1 infected mice retained all three subpopulations of neutrophils in the vagina at 5 d.p.i., in HSV-2 infected mice the presence of cluster 0 was greatly reduced and the bulk of the neutrophils was composed of cluster 2 and 5 (Figure 3C). One major distinguishing characteristic between “homeostatic” cluster 0 and “infection” clusters 2 and 5 was the extent of ISG expression, in which cluster 0 expressed low levels of genes associated with a type I IFN response, even in infected animals, while clusters 2 and 5 expressed high levels of these genes (Figure 3D) (Liberzon et al., 2015). Furthermore, expression of ISGs within clusters 2 and 5 was higher after HSV-2 infection compared to HSV-1 (Figure 3C), which was confirmed by quantitative RT-PCR (qRT-PCR) analysis of select ISGs that are differentially expressed in the vagina at 5 days after HSV-1 or HSV-2 infection (Figure 3 - Supplement 1). qRT-PCR shows that expression of CXCL10 (Figure 3 - Supplement 1A, B) and Gbp2 (Figure 3 - Supplement 1C, D) is increased in HSV-2 infected vaginas compared to HSV-1, while IL-15 is not (Figure 3 - Supplement 1E, F), which supports the accompanying scRNA-seq analysis. While type I IFN was robustly produced early during acute infection after both HSV-1 and HSV-2 infection (Figure 3 - Supplement 2), greater expression of IFN*β* was detected in the vagina after HSV-2 infection compared to HSV-1 at time points corresponding to the onset of genital inflammation (Figure 3E). Thus, during viral infection, distinct neutrophil subsets can be classified by transcriptional profiling, and expression of ISGs suggest that a key difference between a pathogenic and non-pathogenic neutrophil response during viral infection may be sustained IFN signaling.

**Figure 3.**
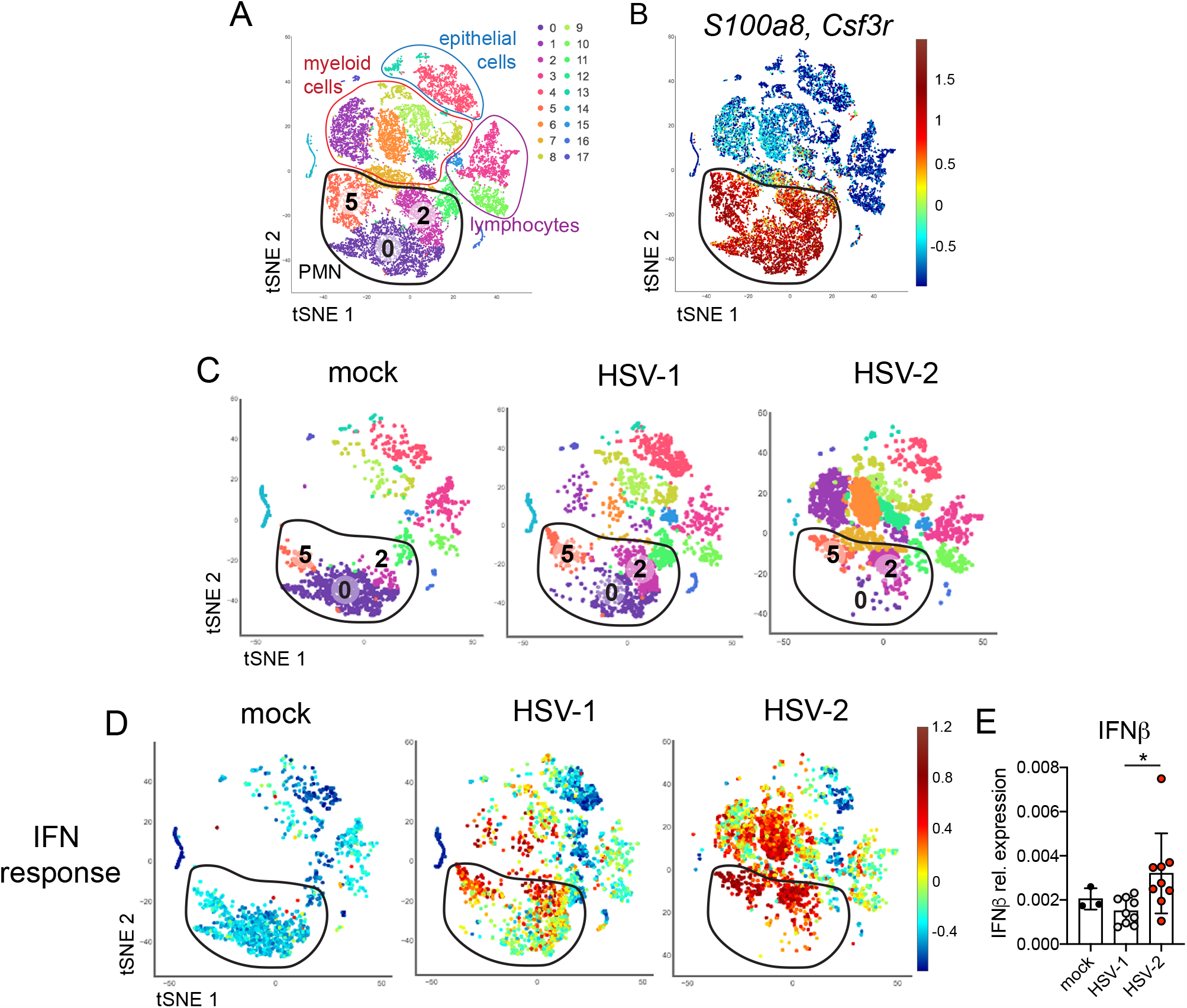
Single cell transcriptome analysis reveals a sustained IFN signature in the neutrophil response against HSV-2. Mice were infected as described in Figure 2. Vaginas were harvested at 5 d.p.i., live cells were flow sorted and subjected to high-throughput scRNA-seq. **A**. A t-Distributed Stochastic Neighbor Embedding (tSNE) visualization of 21,633 cells across all mice resolves 17 distinct clusters in the vaginal tissue. Clusters can be identified as myeloid cells (red border), epithelial cells (blue border) or lymphocytes (purple border). Neutrophils are encircled in black, and contain three distinct clusters (0, 2 and 5). **B**. Neutrophils are defined by high expression of S100A8 and G-CSFR (Csf3r). **C**. tSNE plots of vaginal cell clusters from mock inoculated or HSV-infected mice. Neutrophil populations are circled in black. **D**. Distribution of expression for genes within the Hallmark IFNα Response gene set. **E**. Expression of IFNβ transcripts normalized to RPL13 in mock inoculated (n=3) mice or mice at 5 d.p.i. with HSV-1 (n=9) or HSV-2 (n=9) as measured by qRT-PCR. scRNA-seq in **A-D** was performed once. Data in **E** are pooled from 2 independent experiments. Statistical significance was measured by one-way ANOVA with Tukey’s multiple comparisons test. *p<0.05. Raw values for each biological replicate and specific p values for E are provided in Figure 3 - Source Data.

### Sustained cell-intrinsic type I IFN signaling is required for pathogenic neutrophil responses during HSV-2 infection

We next wanted to test whether type I IFN signaling promoted immunopathology during genital HSV-2 infection. IFNAR1-deficient mice are highly susceptible to HSV regardless of the route of inoculation (Gill, Deacon, Lichty, Mossman, & Ashkar, 2006; Iversen, Ank, Melchjorsen, & Paludan, 2010; Iversen et al., 2016; Reinert et al., 2012; Royer et al., 2019; Svensson, Bellner, Magnusson, & Eriksson, 2007; Wilcox, Folmsbee, Muller, & Longnecker, 2016), and rapidly succumb to infection, mainly due to a loss of viral control. To investigate the temporal effects of type I IFNs in HSV-2 genital disease, we used an antibody against IFNAR1 to block the receptor at different time points after infection (Scott et al., 2018). When mice were injected i.p. with anti-IFNAR1 antibody on the day of HSV-2 inoculation, disease progression was more rapid compared to isotype control treated animals (Figure 4 - Supplement 1A), and mice succumbed to infection at a faster rate (Figure 4 - Supplement 1B), in a manner similar to IFNAR1-deficient mice (Iversen et al., 2010; Iversen et al., 2016; A. J. Lee et al., 2017; Reinert et al., 2012; Wang et al., 2012). Inflammation and rapid disease progression were likely due to significantly elevated viral burden in the anti-IFNAR1 antibody treated mice compared to isotype controls (Figure 4 - Supplement 1C), as HSV is a highly lytic virus that is capable of independently inducing epithelial tissue damage (Horbul, Schmechel, Miller, Rice, & Southern, 2011). To focus on the effects of persistent IFN signaling in the vagina after HSV-2 infection, we also treated mice with a single injection of anti-IFNAR1 antibody or an isotype control at 4 d.p.i.. In stark contrast to early anti-IFNAR1 antibody treatment, one treatment with therapeutic IFNAR1 blockade led to a significant reduction in the severity of inflammation compared to isotype controls (Figure 4A). Histology of vaginal tissues from isotype-treated controls at 6 d.p.i. showed widespread epithelial denuding and immune cell infiltrates within the epithelial layer of the vagina (Figure 4B). In contrast, damage to the vaginal epithelium in anti-IFNAR1 antibody-treated mice appeared to be localized (Figure 4B), similar to neutrophil-depleted mice (Figure 1C). Similarly, the genital skin of isotype control-treated mice displayed signs of severe inflammation and destruction of the epidermis, while the skin structure of anti-IFNAR1 antibody-treated mice was largely intact (Figure 4B). Furthermore, IFNAR1 blockade at 4 d.p.i. had little impact on mucosal viral shedding (Figure 4C). Collectively, these data show that the protective effect of type I IFN on control of genital HSV infection is limited to the early stages of acute infection, and that sustained IFN signaling in the later stages of acute HSV-2 genital infection drive inflammation with minimal effect on viral replication.

**Figure 4.**
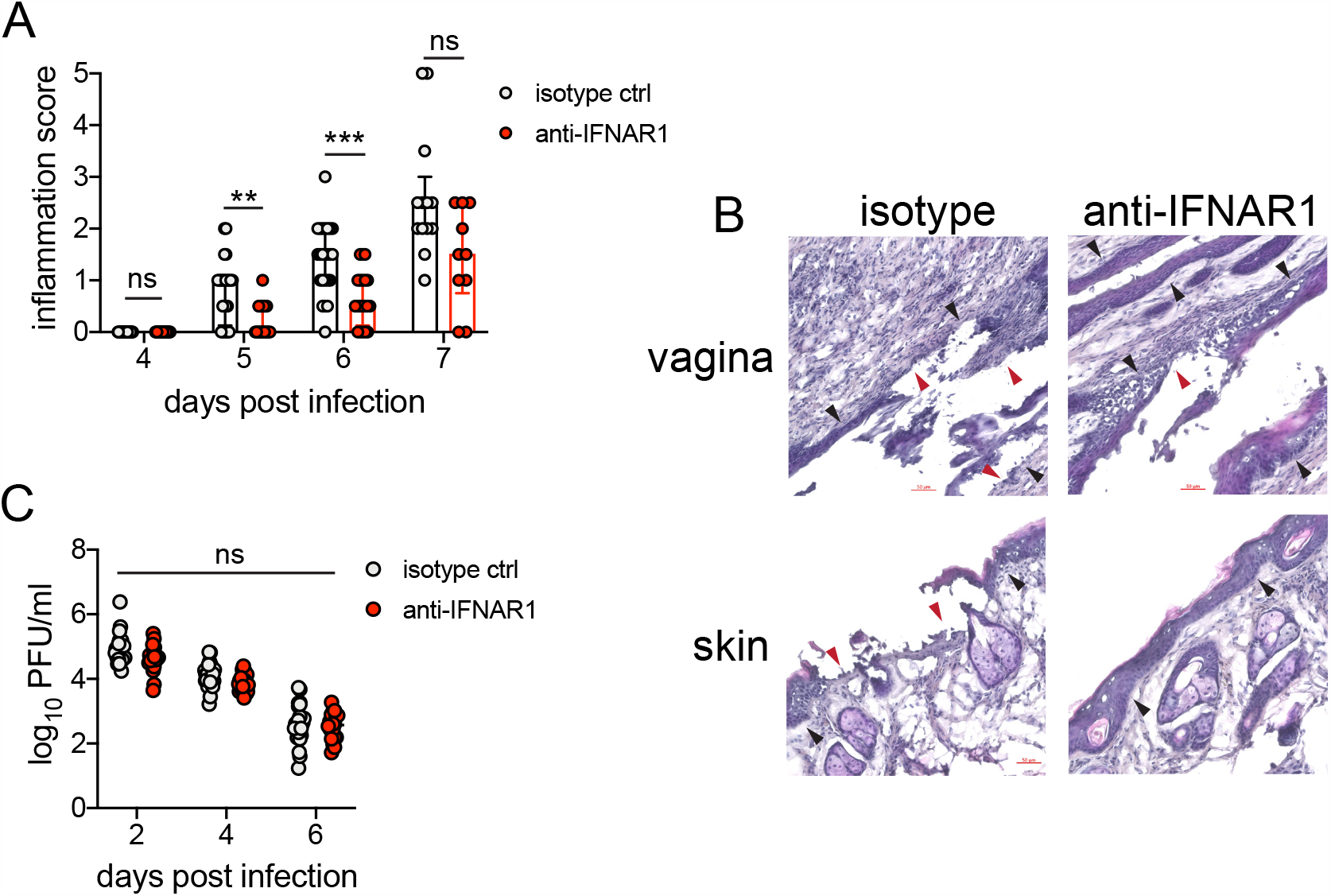
Inhibition of type I IFN signaling during the resolution phase of infection reduces inflammation after HSV-2 infection. Mice were infected as described in Figure 2. At 4 d.p.i., mice were injected i.p. with either 1mg of anti-IFNAR1 antibody (n=10-13) or isotype control (n=7-9) and monitored for disease progression. Mice showing overt signs of genital inflammation at the time of antibody injection (4 d.p.i.) were excluded from the study. **A**. Inflammation scores of antibody-treated mice over the first 7 d.p.i.. **B**. Histology of the vagina (top) or genital skin (bottom) at 6 d.p.i. Red arrows point to areas of epithelial denuding or damage. Black areas denote the basement membrane. **C**. Infectious virus as measured by plaque assay in vaginal washes collected on the indicated days. Data are pooled from (**A, C**) or representative of 3 independent experiments. Bars in **A** show median with interquartile range, bars in **C** show mean. Statistical significance was measured by repeated measures two-way ANOVA with Geisser-Greenhouse correction and Bonferroni’s multiple comparisons test (**A**) or two-way ANOVA with Bonferroni’s multiple comparisons test (**C**). *p<0.05, **p<0.01, ***p<0.005, ns = not significant. Raw values for each biological replicate, epsilon values and specific p values are provided in Figure 4 - Source Data.

Single-cell transcriptional profiling data suggested that type I IFN signaling was highly robust in vaginal neutrophils after HSV-2 infection (Figure 3D). To determine whether intrinsic IFN signaling in neutrophils promoted immunopathology, we deleted the type I IFN receptor from granulocytes by breeding IFNAR1^fl/fl^ x MRP8-Cre mice (IFNAR1 CKO). After confirming that IFNAR1 ablation was limited to the neutrophil population (Figure 5A), IFNAR1 CKO mice and littermate Cre-controls were vaginally infected with HSV-2. Despite differences in IFNAR1 expression, the number of neutrophils recovered from the vaginal lumen was similar between the IFNAR1 CKO mice and their Cre-control littermates (Figure 5B). Strikingly, although the magnitude of the vaginal neutrophil response was similar, we found that the severity of genital inflammation presented by the IFNAR1 CKO mice was significantly reduced compared to the Cre-controls (Figure 5C). As observed after neutrophil depletion, a subset of the IFNAR1 CKO cohort did not develop any signs of inflammation as late as 7 d.p.i. (Figure 5C). Similar to our observations with therapeutic IFNAR1 blockade, IFNAR1 CKO mice exhibited less pathology in both the vagina and genital skin compared to Cre-controls. (Figure 5D). Distinct disease outcomes between the Cre-controls and IFNAR1 CKO mice occurred independently of viral control, as viral load in the mucosa were similar between the two groups (Figure 5E). Together, our data demonstrates that tissue inflammation during HSV-2 infection is largely driven by prolonged type I IFN production, which acts directly upon neutrophils to drive disease.

**Figure 5.**
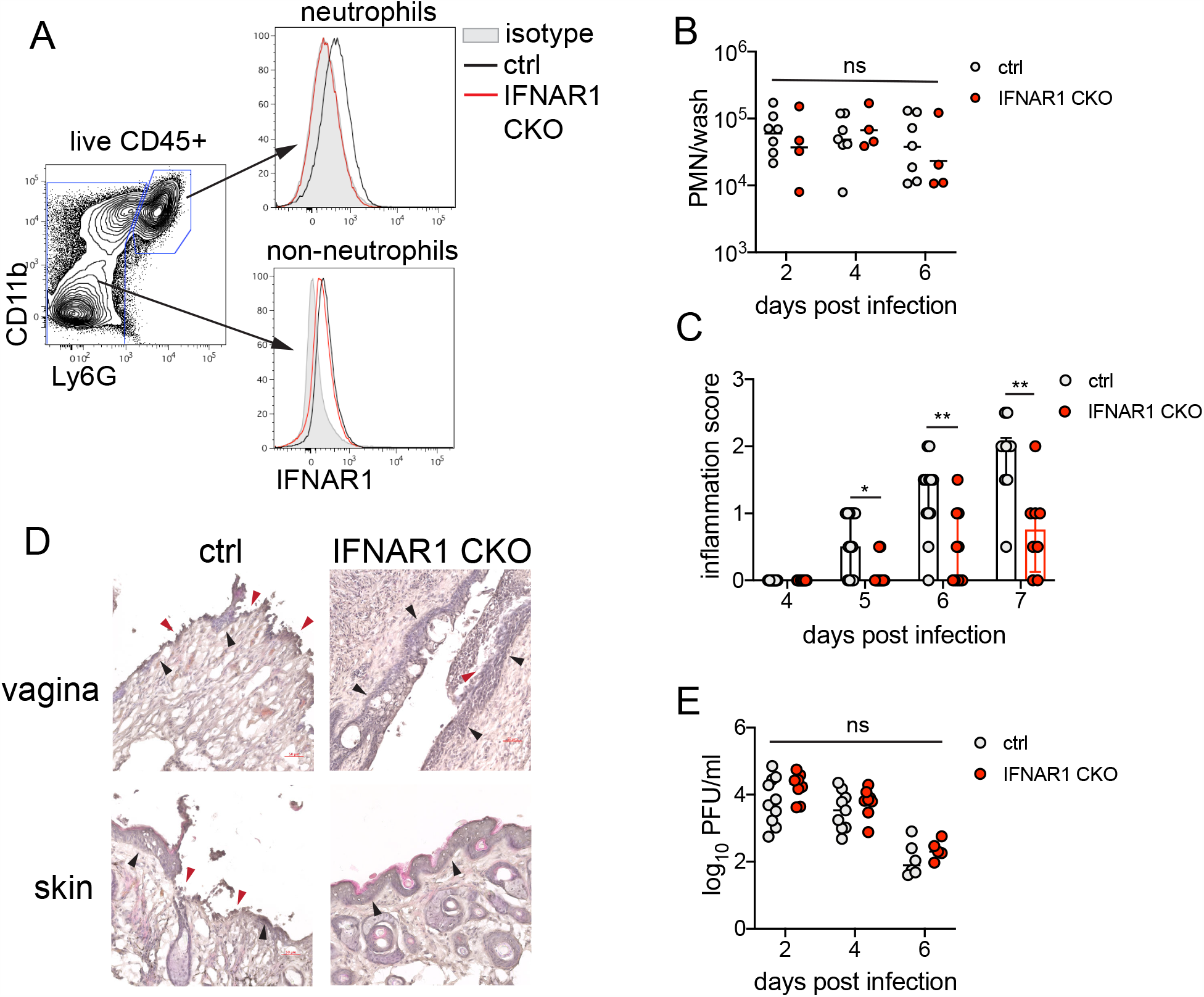
Type I IFN signaling in neutrophils promotes genital inflammation after HSV-2 infection. **A**. IFNAR1 expression on neutrophils and non-neutrophil hematopoietic cells from the bone marrow of naive IFNAR1^fl/fl^ x MRP8-Cre (IFNAR1 CKO) or Cre- littermate controls. Plot is gated on live CD45+ cells. CD11b+Ly6G+ cells are neutrophils, Ly6G-cells are non-neutrophils. Gray histogram shows isotype staining, black open histogram is Cre-control, and red open histogram is IFNAR1 CKO. **B**. Neutrophils were counted by flow cytometry in vaginal washes collected at the indicated days from IFNAR1 CKO (n=4) or Cre-controls (n=7) that were infected with HSV-2 as described in Figure 1. **C**. Inflammation scores for the first 7 d.p.i. of IFNAR1 CKO (n=10-13) or Cre-controls (n=8-11). **D**. Histology on vagina and genital skin at 6 d.p.i.. Red arrows point to areas of epithelial denuding or damage, black arrows denote basement membrane E. Infectious virus as measured by plaque assay from vaginal washes collected on the indicated days from IFNAR1 CKO (n=5-8) or Cre-controls (n=7-10). Data in **C** and **E** are pooled from 3 independent experiments, data in **B** are pooled from 2 independent experiments, and data in **D** are representative of 2 independent experiments. Bars in **C** show median with interquartile range, bars in **B** and **E** show mean. Statistical significance was measured by mixed-effects analysis with (**C**) or without (**B, E**) Geisser-Greenhouse correction and Bonferroni’s multiple comparisons test. *p<0.05, **p<0.01, ns = not significant. Raw values for each biological replicate, epsilon values and specific p values are provided in Figure 2 - Source Data.

### Sustained type I IFN signaling and neutrophils regulate production of pathogenic IL-18 in the vagina during HSV-2 infection

Type I IFN stimulation of neutrophils can upregulate ISGs as well as several pro-inflammatory cytokines (Galani et al., 2017). To determine whether type I IFN was driving disease by shaping the cytokine milieu within the vagina, we first measured several pro-inflammatory cytokines in the vagina at 5 d.p.i., in the presence or absence of neutrophils. The production of inflammatory cytokines such as IL-6 (Figure 6 - Supplement 1A), IL-1*β* (Figure 6 - Supplement 1B) or TNF (Figure 6 - Supplement 1C), all of which have been associated with genital inflammation and HSV-2 infection in humans (Gosmann et al., 2017; Masson et al., 2014; Murphy & Mitchell, 2016), was similar between both neutrophil-depleted and control groups. Production of IFN*γ* (Figure 6 - Supplement 1D) as well as IL-12p70 (Figure 6 - Supplement 1E), both cytokines associated with a type I immune response and important for HSV control, were similar between the neutrophil-depleted and control groups. However, when we measured IL-18, an IL-1 family cytokine that primarily known for mediating innate defense (Harandi, Svennerholm, Holmgren, & Eriksson, 2001) and for promoting IFN*γ* production from NK cells during genital HSV-2 infection (A. J. Lee et al., 2017), we detected a notable difference between neutrophil-depleted and control mice (Figure 6A), suggesting an unexpected role for this cytokine in driving disease during HSV-2 infection.

**Figure 6.**
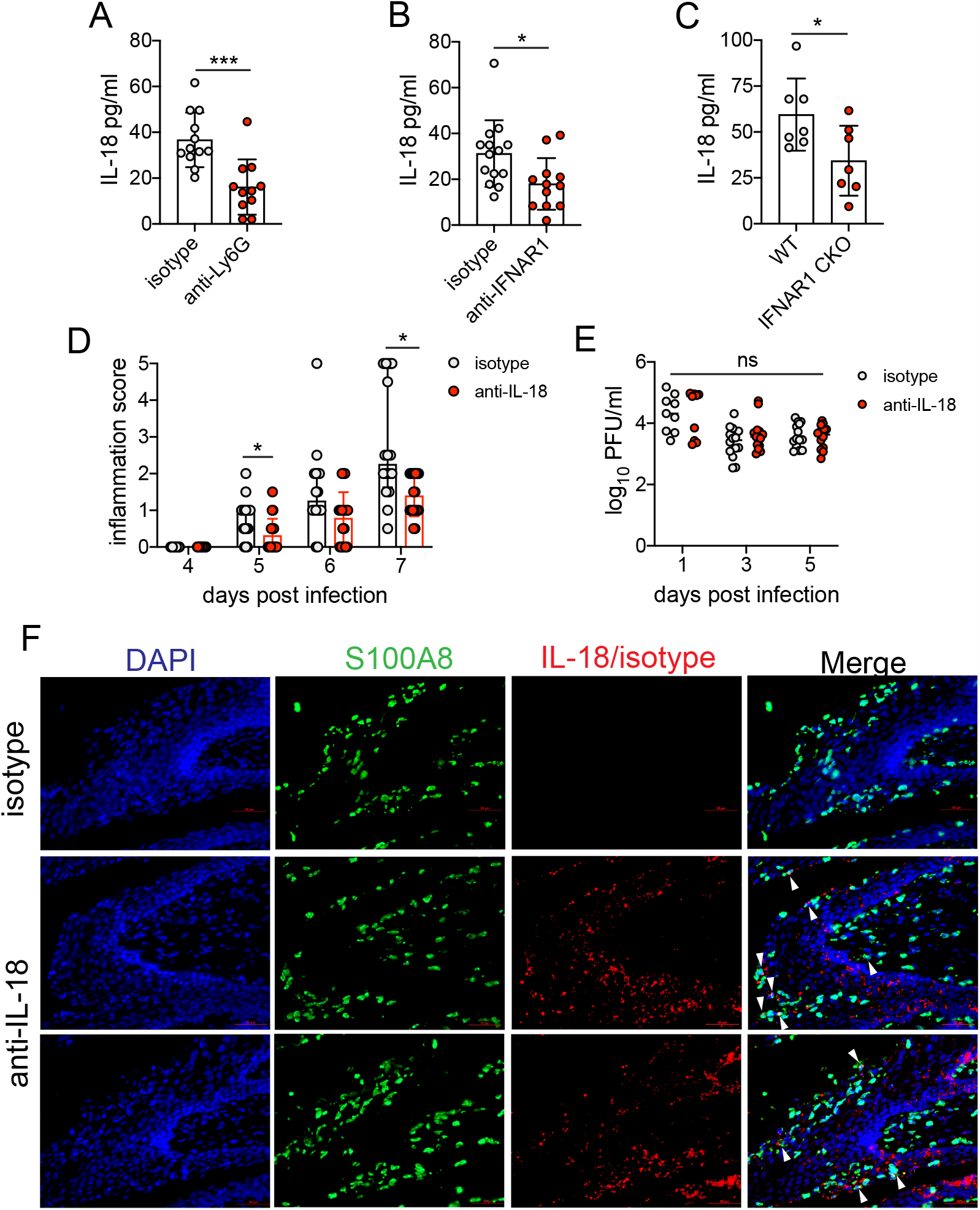
Sustained type I IFN signaling and neutrophils regulate pathogenic IL-18 levels in the vagina. C57BL/6 mice were treated with anti-Ly6G (n=11) or isotype control (n=12) as described in Figure 1 (**A**), therapeutically treated with anti-IFNAR1 (n=12) or isotype control (n=14) as described in Figure 5 (**B**) and infected with HSV-2, or IFNAR1 CKO (n=7) and Cre-controls (n=7) were infected with HSV-2 as described in Figure 5 (**C**). **A-C**. Vaginal IL-18 levels were measured by ELISA in washes collected at 5 d.p.i.. 100μg of anti-IL-18 neutralizing antibody (n=18) or isotype control (n=16) was administered ivag at 3, 4 and 5 d.p.i.. **D**. Inflammation scores of antibody-treated mice over the first 7 d.p.i.. **E**. Infectious virus as measured by plaque assay in vaginal washes collected on the indicated days (n=9-14). **F**. Immunofluorescent staining of vaginal tissues collected at 6 d.p.i.. Green shows S100A8 (neutrophil), red shows IL-18 or isotype, blue is DAPI. Top row shows tissues probed with isotype control, and bottom two rows show two representative images of tissues probed with anti-IL-18 antibody. White arrows point to IL-18+ neutrophils. Data in **A-C, D** and **E** are pooled from 4 independent experiments for each experimental setup. Data in **F** is representative of 2 independent experiments. Bars in **A-C** show mean and SD, bars in **D** show median with interquartile range, and bars in E show mean. Statistical significance was measured by unpaired t-test (**A-C**), repeated measures two-way ANOVA with (**D**) Geisser-Greenhouse correction and Bonferroni’s multiple comparisons test, or mixed-effects analysis with Bonferroni’s multiple comparisons test (**D**). *p<0.05, ***p<0.005, ns = not significant. Raw values for each biological replicate, epsilon values and specific p values are provided in Figure 2 - Source Data.

To determine whether type I IFN signaling regulated IL-18 production in the vagina, we assessed IL-18 levels in the vaginal lumen after therapeutic antibody-mediated IFNAR1 blockade. At 5 d.p.i., similarly to neutrophil-depleted mice, we found that IL-18 levels were markedly reduced (Figure 6B). Importantly, measurement of IL-18 in the vagina of IFNAR1 CKO at 5 d.p.i. also revealed a significant decrease in cytokine levels compared to littermate controls (Figure 6C). To determine whether IL-18 was playing a key role in driving immunopathology during genital HSV-2 infection, we therapeutically administered an IL-18 neutralizing antibody to HSV-2 infected animals starting at 3 d.p.i.. Remarkably, neutralization of IL-18 led to a considerable reduction in disease severity (Figure 6D), without any impact on viral control (Figure 6E). To determine the source of pathogenic IL-18 in the vagina, we probed vaginal tissues for the neutrophil marker S100A8 and IL-18 at 6 d.p.i. (Figure 6F). Detection of IL-18 and S100A8 around a single nucleus demonstrated that neutrophils could be a source of IL-18 during vaginal HSV-2 infection (Figure 6F). However, we also identified IL-18-reactive cells that were negative for S100A8 but in close proximity to neutrophils (Figure 6F), suggesting the potential for multiple cellular sources of IL-18. Thus, our data demonstrate that sustained type I IFN signaling in neutrophils leads to the production of vaginal IL-18, and reveal IL-18 to be a novel regulator of disease after HSV-2 infection.

## DISCUSSION

In this study, we evaluated drivers of a pathogenic neutrophil response using a mouse model for an important human infection. We found that neutrophils promote genital inflammation and do not impact antiviral activity after genital HSV-2 infection, suggesting that the neutrophil response is primarily immunopathogenic. Depletion of neutrophils led to a significant decrease in disease severity without affecting recruitment of other immune cells or the production of common pro-inflammatory cytokines, and deficiency in genes controlling neutrophil effector functions such as ROS production and NET formation had little impact on progression of disease. Comparative analysis of single-cell transcriptional profiles revealed a strong type I IFN signature that was sustained in neutrophils responding to a highly inflammatory genital HSV-2 infection but not a less inflammatory HSV-1 infection. In contrast to antibody-mediated blockade of IFNAR1 at the time of infection, which led to significantly worse disease outcomes, IFNAR1 blockade just prior to the resolution phase of acute mucosal infection significantly delayed the progression of genital inflammation. Importantly, neutrophil-specific deficiency of IFNAR1 markedly reduced the severity of genital disease after HSV-2 infection, suggesting that persistent IFN signaling drove disease primary by acting on neutrophils. Ultimately, this sustained type I IFN signaling in neutrophils promoted the production of pro-inflammatory IL-18, and therapeutic neutralization of this cytokine also ameliorated disease. Together, our results suggest an axis of type I IFNs, neutrophils, and IL-18 as key drivers of genital disease in a mouse model of HSV-2 infection, and that sustained type I IFN signaling is a key factor in distinguishing between pathogenic and non-pathogenic neutrophil responses during mucosal viral infection.

Type I IFNs are a frontline of defense against viral infection, but models of chronic viral infection, including lymphocytic choriomeningitis virus (LCMV) (Teijaro et al., 2013; Wilson et al., 2013), human immunodeficiency virus (HIV) (Meier et al., 2009; Rotger et al., 2010; Sedaghat et al., 2008; Taleb et al., 2017) and simian immunodeficiency virus (SIV) (Harris et al., 2010; Jacquelin et al., 2009), reveal the detrimental effect of overexuberant or sustained type I IFN signaling. Notably, prolonged IFN signaling during chronic viral infection can promote immunosuppression through multiple cellular and molecular mechanisms, and deletion or blockade of IFNAR1 during chronic LCMV infection can alleviate immunosuppression and enhance long-term viral control (Cheng et al., 2017; Taleb et al., 2017; Teijaro et al., 2013; Wilson et al., 2013). However, unlike the LCMV model, early blockade of type I IFN signaling led to more severe disease and a complete loss of viral control after HSV-2 infection, similar to infections performed on an IFNAR1-deficient genetic background (Iversen et al., 2010; Iversen et al., 2015; A. J. Lee et al., 2017; Leib et al., 1999; Reinert et al., 2012; Wang et al., 2012) and indicating an early antiviral role (A. J. Lee et al., 2017; Luker, Prior, Song, Pica, & Leib, 2003). Rather, only inhibition of sustained IFN signaling led to diminished disease with minimal impact on viral control, thus revealing distinct antiviral and immunopathological effects of type I IFN signaling that are temporally regulated and had heretofore been unappreciated during HSV-2 infection. The source of sustained type I IFN production that promotes immunopathology after genital HSV-2 infection is currently unknown. HSV encodes numerous proteins that can suppress type I IFN production and regulate the signaling pathways (Christensen et al., 2016; Lin & Zheng, 2019; Melroe, DeLuca, & Knipe, 2004), suggesting that production of type I IFN likely occurs from a cell type that is not directly infected. While plasmacytoid dendritic cells (pDC) are known as robust producers of type I IFN, previous studies have demonstrated a limited role for pDCs in genital HSV-2 infection (Swiecki, Wang, Gilfillan, & Colonna, 2013), indicating that there may be an alternative source of type I IFN, such as conventional DCs (Wilson et al., 2013).

In humans, type I IFN can be detected at active lesion sites during recurrent episodes (Peng et al., 2009; Roychoudhury et al., 2020), although levels do not correlate with restriction of viral replication (Roychoudhury et al., 2020). This raises the possibility that type I IFN induction may not be antiviral and could contribute to ulcer formation, although this hypothesis has yet to be tested. Human neutrophils from females are also reported to be hyper-responsive to type I IFNs (Gupta et al., 2020). Although clinical disease recurrence rates between men and women with genital herpes are similar (Wald et al., 2002), differences in neutrophil sensitivity to type I IFN may have implications for sex-dependent mechanisms of ulcer development.

Type I IFN signaling orchestrates a network of ISGs that limit viral infection using a wide variety of mechanisms. In vitro stimulation of neutrophils with type I IFN leads to the upregulation of many common ISGs as well as inflammatory genes, including IL-18 (Galani et al., 2017). Importantly, it has been suggested that type I IFN can differentially regulate expression IL-18 and IL-1*β*, another IL-1 family cytokine that depends on caspase-mediated cleavage for activation (Zhu & Kanneganti, 2017). It is currently unclear whether neutrophils are directly producing this cytokine in our model of infection and whether IL-18 production is dependent on the classical inflammasome-dependent route. As HSV also encodes proteins that can inhibit inflammasome activity, including VP22 (Maruzuru et al., 2018), one possibility is that that the direct source of IL-18 is a cell type that is not productively infected with HSV, such as neutrophils. Alternatively, neutrophil proteases released in the extracellular space have been reported to cleave and activate proIL-1 cytokines that are secreted by other cells in a caspase-1-independent manner (Clancy et al., 2018; Robertson et al., 2006; Sugawara et al., 2001), suggesting a mechanism by which neutrophils may modulate IL-18 levels without directly producing the cytokine themselves. Our data show that along with neutrophils, IL-18 could be detected in the epithelium in cells that are in close proximity to infiltrating neutrophils, indicating that there may be multiple sources and mechanisms by which pathogenic IL-18 is produced during HSV-2 infection. Understanding these mechanisms could further reveal novel, specific targets for therapeutics aimed at reducing inflammation during genital herpes, and as such, are currently under investigation.

During HSV-2 vaginal infection, IL-18 stimulates NK cells to mediate rapid antiviral IFN*γ* production upon infection (A. J. Lee et al., 2017), and previously thought to be important for orchestrating a protective innate immune response. Accordingly, mice deficient in IL-18 are more susceptible to HSV-2 infection (Harandi et al., 2001), as well as infection with HSV-1 through multiple routes of inoculation (Fujioka et al., 1999; Reading et al., 2007), presumably due to dysregulation of innate IFN*γ* production and loss of viral control. Our study reveals a novel aspect of IL-18 biology during HSV-2 infection, and that like type I IFN signaling, there may be a temporal component to the effects of IL-18 during HSV-2 infection. Therapeutic neutralization of IL-18 did not alter viral titers in our model, suggesting that IL-18 does not have an impact on T cell-dependent IFN*γ* production (Milligan & Bernstein, 1997; Nakanishi, Lu, Gerard, & Iwasaki, 2009) or direct antiviral activity. As previous studies have shown that IL-18 is also dispensable for stimulating IFN*γ* from adaptive memory immune responses (Harandi et al., 2001), IL-18 may be an attractive target for therapeutics aiming to reduce inflammation during genital herpes. Currently, the mechanism by which IL-18 promotes disease during genital HSV-2 infection is unknown. In the gut, the role of IL-18 is balanced between protection and pathology, depending on the source of IL-18, the model of disease and the responsive cell type (Jarret et al., 2020; Nowarski et al., 2015). The role of IL-18 during HSV-2 infection appears to be similarly complex, and further study will be required to identify the compartment on which IL-18 acts and the downstream effects of IL-18 signaling. Additionally, while our results demonstrate an important role for IL-18, the reduction in disease severity is not as profound as when type I IFN signaling is inhibited in our HSV-2 model of infection. Considering the complex response elicited by type I IFN, our data allude to the possibility of other IL-18-independent, IFN-dependent mechanisms that promote genital inflammation that have yet to be elucidated.

Synergistic effects of cytokine signaling have been reported to be important for maximizing cellular responses to infection through the upregulation of cooperative or independent molecular programs (Bartee & McFadden, 2013) or through the cross-regulation of receptor signaling pathways (Ivashkiv & Donlin, 2014). Along with sustained type I IFN signaling during HSV-2, the vaginal cytokine milieu also changes during the course of genital HSV-2 infection. Whether the ISG profile induced in neutrophils by type I IFN signaling changes over the course of acute HSV-2 infection, and whether this profile is affected by other cytokines, is unknown. Upon activation, the activity of neutrophils responding to infection can be modulated strongly by multiple interferons (IFNs), in a variety of tissues. Type I (Ank et al., 2008), type II (Iijima et al., 2008; A. G. Lee et al., 2020) and type III IFNs (Ank et al., 2008; A. G. Lee et al., 2020) are all robustly produced during HSV-2 infection. While type I and type II IFNs are crucial for control of HSV-2 replication, endogenous type III IFNs do not appear to affect either disease severity or viral control, although exogenous application of type III IFNs can reduce viral burden (Ank et al., 2008; Ank et al., 2006). Expression of the type III IFN receptor, IFNLR, is limited to very few cell types, including neutrophils and epithelial cells (Blazek et al., 2015; Mahlakõiv, Hernandez, Gronke, Diefenbach, & Staeheli, 2015; Sommereyns, Paul, Staeheli, & Michiels, 2008). As epithelial cells are a major target for HSV-2 replication, dissecting the action of type III IFNs within the neutrophil and epithelial cell compartments may reveal a more detailed picture of the role type III IFNs play. The impact of simultaneous type I, II and III IFN signaling on neutrophil function is currently unclear and due to the importance of these molecules, as well as PRRs, in controlling infection, cell-specific modifications of receptor expression will be required for further investigation.

Beyond their inflammatory role during HSV-2 infection, this study and others (S. Li et al., 2018; Milligan, 1999) have demonstrated that neutrophils can be found in the vagina at steady state. Detection of neutrophils in washes collected from the vaginal lumen suggest that the neutrophils are actively extravasating, migrating through the lamina propria and through the epithelial barrier. Previous studies have indicated that in mice, recruitment of neutrophils into the vagina is regulated by hormone-dependent expression of chemokines, including CXCR2 ligands, in the tissue (Lasarte et al., 2016). In humans, the number of neutrophils in the fluctuates with the hormone cycle in the upper reproductive tract, but reportedly remains stable in the lower tract (Wira, Rodriguez-Garcia, & Patel, 2015). It is unclear why neutrophils actively patrol the vagina even in the absence of infection, what consequences their immunosurveillance has on the normal physiology of the reproductive tract, and how the tissue microenvironment affects the biology of the neutrophils themselves. In the oral mucosa, constant recruitment of neutrophils into the tissue helps support the maintenance of a healthy microbiome (Uriarte, Edmisson, & Jimenez-Flores, 2016), while molecular products derived from the microbiome such as peptidoglycan can support basal activity of neutrophils (Clarke et al., 2010). Whether similar interactions between the vaginal neutrophils and the vaginal microbiome also occur is unknown. Furthermore, due to the lack of tractable animal models, the role of neutrophils in other common non-viral STIs is unclear, although neutrophils appear to play a similar pathological role in the upper reproductive tract after Chlamydia infection (Lijek, Helble, Olive, Seiger, & Starnbach, 2018). In summary, our study reveals that pathology during genital HSV-2 infection is driven at least in part by a robust neutrophil response, and lays the foundation for identifying the host pathways that may be targeted to help ameliorate disease.

## ACKNOWLEDGEMENTS

We thank Rachel Idol, Antonina Akk and Celeste Cummings for technical assistance on assays used in this study. This work was supported by grants from the NIH (HS: R01 AI134962) and Children’s Discovery Institute (RAC). T.J.L was supported by funding for the Training Program in Immunology from the NIH (T32 AI007163).

## Author contributions

T.J.L, Y.S.L., A.C., M.R.F., J.M.S., A.N.O., X.J., and H.S. designed and conducted experiments, acquired data and analyzed data. R.A.C., C.T.N.P, and M.C.D. aided in the design of experiments. P.S.A. and M.A. analyzed data. H.S. and Y.S.L. wrote the manuscript, and all authors edited the final version of the paper.

### SOURCE DATA

Source data files have been provided for all figures and figure supplements.

## METHODS

### Mice

Six-week old female C57BL/6J mice were purchased from Jackson Laboratories and rested for at least one week and infected at a minimum of seven weeks of age. Ncf2 KO mice and controls were provided by M.C. Dinauer (Washington University, St Louis) and generated as previously described (Jacob et al., 2017). Stim1^fl/fl^ x Stim2^fl/fl^ x MRP8-Cre mice were provided by G.A. Clemens (Washington University, St Louis) and were generated as previously described (Clemens et al., 2017). IFNAR1^fl/fl^ mice (Ifnar1^tm1Uka^) were a gift from H.W. Virgin (Kamphuis, Junt, Waibler, Forster, & Kalinke, 2006; Nice et al., 2016). PAD4^fl/fl^ mice (B6(Cg)-Padi4^*tm1*.*2Kmow*^/J) and MRP8-Cre (B6.Cg-Tg(S100A8-cre,-EGFP)1Ilw/J) were obtained from Jackson Laboratories and bred at Washington University School of Medicine. Crelittermates generated from breeding pairs were used as controls. All mice were maintained on a 12 hour light/dark cycle with unlimited access to food and water. This study was carried out in accordance with the recommendations in the Guide for the Car and Use of Laboratory Animals of the National Institutes of Health.

### Ethics statement

The protocols were approved by the Institutional Animal Care and use Committee (IACUC) at the Washington University School of Medicine (Assurance number A3381-01). All experiments were performed under biosafety level 2 (A-BSL2) containment and all efforts were made to minimize animal suffering.

### Cell lines and primary cells

Vero Cells (African green monkey kidney epithelial cells, ATCC) were cultured in Dulbeco’s Modified Eagle Medium (Gibco) containing 1% fetal bovine serum (FBS, Corning) and maintained at 37°C with 5% CO2. Primary neutrophils were isolated from the bone marrow (BM) of naive female C57BL/6J mice. A Histopaque gradient was used to isolate primary neutrophils for Reactive Oxygen Species (ROS) assays, while a Percoll gradient was used for NET assays. For Histopaque isolation: 3ml of Histopaque 1119 (Sigma-Aldrich) was overlaid with 3ml of Histopaque 1077 (Sigma-Aldrich). A single cell suspension of isolated BM cells in 1ml of PBS was layered over the Histopaque gradient. Cells were centrifuged for 30 minutes at room temperature (RT), and neutrophils were collected from the bottom interface. For Percoll isolation: BM cells were resuspended in HBSS (Gibco) with 20mM HEPES (Gibco) and layered over 6ml of 62% Percoll solution (GE Healthcare). Cells were centrifuged for 30 minutes at RT, and neutrophils were collected from the bottom of the tube. All tissue culture experiments were performed under BSL2 containment.

### Viruses and virus quantification

WT HSV-2 186 syn+ (Spang, Godowski, & Knipe, 1983) and HSV-1 McKrae (Williams, Nesburn, & Kaufman, 1965) was propagated and titered on Vero cells as previously described (A. G. Lee et al., 2020). Briefly, for propagation of virus stocks, Vero cells were plated in T150 tissue culture flasks, inoculated at 0.01 MOI at 80% confluence and incubated at 37°C. Infected cells were harvested 2-3 days after infection, resuspended in equal volumes of virus supernatant and twice-autoclaved milk, and sonicated. Lysed cells were aliquoted and used as viral stock. To titer, Vero cells were plated in 6-well plates and inoculated with 10-fold serial dilutions of stock virus. After inoculation, overlay media with 20*μ*g/ml human IgG was added to each well and plates were incubated at 37°C for 2-3 days. To count, Vero cells were stained with 0.1% crystal violet. All tissue culture experiments were performed under BSL2 containment. For titration of virus in the vaginal lumen, 50ul washes with sterile PBS were collected using a pipette and a sterile calginate swab, and diluted in 950ul of ABC buffer (0.5mM CaCl2, 0.5mM MgCl2, 1% glucose, 1% fetal bovine serum (FBS) in sterile PBS). 10-fold serial dilution of vaginal washes were titered by plaque assays won Vero cells (A. G. Lee et al., 2020).

### Mouse infection studies

All mice were in injected subcutaneously in the neck ruff once with 2mg of DMPA (Depo-Provera, Pfizer) 5-7 days prior to virus inoculation. For experiments in which neutrophils were depleted, mice were intraperitoneally (i.p.) injected once with 500*μ*g of anti-Ly6G (clone 1A8) or rat IgG2a isotype control (anti-trinitrophenol+KLH) (Leinco Technologies) diluted in sterile phosphate buffered saline (PBS, Sigma-Aldrich) 1 day prior to inoculation. For experiments in which IFNAR blockade was conducted, mice were i.p. injected once with 1mg of anti-IFNAR1 (clone MAR1-5A3) or mouse IgG1 isotype control (clone HKSP) (Leinco Technologies) on either the day of inoculation (“early”) or at 4 d.p.i. (“late”). For “late” treatments, only mice without overt signs of genital inflammation were chosen for antibody injection in both anti-IFNAR and isotype control groups to avoid biasing of results. For experiments in which IL-18 was neutralized, mice were treated intravaginally with 100*μ*g of anti-IL-18 antibody (clone YIGIF74-1G7) or rat IgG2a isotype control (clone 2A3) (BioXCell) on days 3-5 after infection. Selection of mice for isotype control or experimental antibody treatment was random. For intravaginal inoculation, a sterile calginate swab (McKesson) moistened with sterile PBS was used to gently disrupt mucous from the vaginal cavity. Stock virus was diluted in sterile PBS and either 5000 PFU or 10^4^ PFU virus was delivered into the vaginal cavity via pipette tip in a 10*μ*l volume. Mice were weighed and monitored for signs of disease for 1 week following infection and monitored for survival for 2 weeks. Genital inflammation was scored as follows: 0 - no inflammation, 1 - mild redness and swelling around the vaginal opening, 2 - fur loss and visible ulceration, 3 - severe ulceration and mild signs of sickness behavior (lack of grooming), 4 - hindlimb paralysis, and 5 - moribund.

### Vaginal tissue processing

All tissues were harvested from animals sedated with ketamine and xylazine and thoroughly perfused with a minimum of 15ml of PBS. Vaginas were processed as follows: tissue was cut into pieces and digested for 15 minutes in a shaking water bath held at 37°C in a 0.5mg/ml solution of Dispase II (Roche) in PBS. Tissues were then transferred to a solution of 0.5mg/ml Collagenase D (Roche) and 15*μ*g/ml DNase I (Roche) in RPMI media (Gibco) supplemented with 10% fetal bovine serum (FBS, Corning) and 1% pen/strep (Gibco) and digested for 25 minutes in a shaking water bath held at 37°C. 50*μ*l of sterile EDTA was added to each sample and incubated at 37°C for another 5 minutes. Tissue were then mechanically disrupted through a 70-um cell strainer into a single cell suspension using a 3ml syringe plunger. Tissues were washed with RPMI media with 1% FBS, centrifuged, and resuspended in 200*μ*l RPRM with 1% FBS and 1% pen/strep.

### Flow cytometry

Single cell suspensions from vaginal tissues, or luminal cells collected in vaginal washes were plated in 96-well plates and incubated with Live/Dead Fixable Aqua Dead Cell Stain kit (Molecular Probes) for 15 minutes at room temperature (RT) in the dark. Cells were then incubated with Fc block (anti-CD16/32, Biolegend) for 15 minutes at RT in the dark. Surface staining was performed in FACS buffer (1% FBS and 0.02% sodium azide in PBS) on ice and in the dark using the following antibodies: CD3 (clone 145-2C11), CD4 (clone GK1.5), CD8a (clone 53-6.7), CD11b (clone M1/70), CD45 (clone 30-F11), Gr-1 (clone RB6-8C5), Ly6C (clone HK1.4), Ly6G (clone 1A8), and NK1.1 (clone PK136). All antibodies were purchased from Biolegend. For surface staining of IFNAR1, cells were incubated with an anti-IFNAR1 antibody or a mouse IgG1 isotype control (Leinco Technologies) for 20 minutes at 37°C. Cells were washed and then surface staining of other markers proceeded as described above. Cell counts were performed by adding Precision Count Beads (Biolegend) to samples prior to flow cytometric acquisition. Samples were acquired on an LSR Fortessa (BD Biosciences) and analyzed by FlowJo (Treestar).

### Tissue immunofluorescent (IF) staining and immunohistochemistry (IHC)

All tissues were harvested from animals sedated with ketamine and xylazine and thoroughly perfused with a minimum of 15ml of PBS, followed by 15ml of PLP fixative (0.01M NaIO4, 0.075M lysine, 0.0375M sodium phosphate, 2% paraformaldehyde (PFA)) for IF or 4% PFA for IHC. Tissues were cryoprotected in 30% sucrose, frozen in OCT medium (Sakura) and cut into 7um sections. Cryosections were blocked 5% bovine serum albumin (BSA), 5% goat serum (Jackson Immunoresearch) and 0.1% Triton-X in PBS for 1 hour at RT. HSV antigens were detected with a rabbit anti-HSV primary antibody (Dako), incubated overnight at 4°C, washed in PBS and incubated for 1 hour at RT with a goat anti-rabbit IgG conjugated to AlexaFluor 488 (Life Technologies). S100A8 was detected with a rat anti-mouse S100A8 primary antibody (clone 63N13G5, Novus Biologicals) and a goat anti-rat IgG conjugated to AlexaFluor 568 (Life Technologies) in a similar manner. IL-18 was detected using a biotinylated rat anti-mouse IL-18 primary antibody (clone 93-10C, MBL International). Cryosections were blocked as described above, and then treated with the Avidin/Biotin Blocking Kit (Vector Laboratories) according to manufacturer’s protocol. Endogenous peroxidases were quenched with a 2 % hydrogen peroxide solution. Anti-mouse IL-18 or a rat IgG1 isotype control were incubated overnight at 4°C. The AlexaFluor 647 Tyramide Signal Amplification kit (Invitrogen) was used to visualize IL-18 and used according to manufacturer’s protocol. DNA was visualized with 4′,6-diamidino-2-phenylindole (DAPI) (Life Technologies). Sections were imaged with a Zeiss Cell Observer inverted microscope using a 40x objective, acquired with Zen software and image brightness was adjusted using Photoshop (Adobe). For IHC, sections were probed with an anti-HSV antibody incubated overnight at 4°C (Dako), a donkey anti-rabbit IgG-HRP antibody (Jackson Immunoresearch) for 1 hr at RT and then enzymatically visualized by 3,3’-diaminobenzidine (DAB) enzyme reaction (Sigma-Aldrich). Sections were counterstained with hematoxylin and eosin and images were captured using Zeiss ZEN software on a Zeiss Cell Observer inverted microscope with an Axiocam dual B/W and color camera with a 20x objective. Image brightness was adjusted using Photoshop (Adobe) and merged with Image J64 (NIH).

### RNA extraction and quantitative reverse transcription (RT)-PCR

Harvested tissues were homogenized in RLT buffer (RNeasy Kit, Qiagen) with approximately 100*μ*l of sterile 1.0mm zirconia/silica beads (Biospec Products) in a bead beater. Homogenized tissue samples were processed according to manufacturer’s protocol using the RNeasy Mini Kit (Qiagen) and RNA quality and quantity was assessed on a Nanodrop (ThermoFisher). qRT-PCR was performed in 10*μ*l reactions using the iTaq Universal SYBR Green One-Step kit (Biorad) according to manufacturer’s protocol.

### Single cell RNA-sequencing preparation

Single cell suspensions from digested vaginas were stained with Live/Dead Fixable Aqua Dead Cell Stain kit (Molecular Probes) for 15 minutes at room temperature (RT) in the dark. Live cells were sorted on BD FacsAria II housed in a BSL2 biosafety cabinet. A minimum of 16,000 cells were resuspended in PBS with 2% FBS and 0.2U/*μ*l RNase inhibitor at a concentration of 800-1400 cells/*μ*l, submitted to McDonnell Genome Institute and prepared for droplet-based 3’ end single-cell RNA sequencing using the Chromium 3’ v3 single cell reagent kit per manufacturer’s protocol (10x Genomics). Library sequencing was performed on a NovaSeq S4 (Illumina).

### Cytokine measurement

For cytokine analysis by Bio-Plex Pro Mouse Cytokine 23-Plex Immunoassay (Bio-rad): 2 x 50*μ*l washes with sterile PBS were collected from the vaginal lumen using a pipette. Samples were centrifuged for 3 minutes at 13000*g to remove mucous and cells, and supernatants were added to 200*μ*l of ABC buffer. The assay was performed according to manufacturer instructions, and plates were read on a Luminex Bioplex 100 system (Biorad). For measurement of IL-18, 2 x 50*μ*l washes with sterile PBS were collected from the vaginal lumen and centrifuged to remove mucous and cells. IL-18 was measured using the mouse IL-18 ELISA kit (MBL International) according to manufacturer’s instructions at half-volumes.

### In vitro neutrophil stimulation

To measure ROS production, isolated neutrophils were stimulated with heat-killed HSV-2 (56°C for 30 minutes) at an MOI of 5 for 16 hours at 37°C. ROS levels were quantified using DCFDA Cellular ROS Detection Assay kit (Abcam) according to manufacturer protocol. Fluorescence levels were measured by flow cytometry. To induce NET formation, neutrophils were stimulated with heat-killed HSV-2 at an MOI if 1 for 4 hours at 37°C. Cells were fixed with 8% PFA overnight and probed with a polyclonal rabbit antibody against mouse citrullinated histone H3 (Abcam) for 1 hour at RT in 1% BSA and 0.1% Triton-X for 1 hour at RT, a goat anti-rabbit antibody conjugated to AlexaFluor 488 (Life Technologies) for 1 hour at RT and DAPI diluted in PBS for 6 minutes at RT. Cells were imaged with a Zeiss Cell Observer inverted microscope using a 63x objective and image brightness was adjusted using Photoshop (Adobe).

### Single cell RNA-sequencing analysis

#### Processing data with Seurat package

The Seurat package in R was used for analysis (Butler, Hoffman, Smibert, Papalexi, & Satija, 2018). Cells with mitochondrial content greater than 5% were removed. The initial analysis of the data revealed three clusters of the cells that had extremely low levels of detected genes (i.e. <500) which were then filtered out as non-viable cells. Remaining cells were used for downstream analysis, resulting in the 6,507 cells per sample that passed quality control (QC) and filtering. Filtered data were normalized using a scaling factor of 10,000, and nUMI was regressed with a negative binomial model.

#### Normalization and Feature Selection

After the data filtration, data were normalized using a scaling factor of 10,000 and log transformed. The highly variable genes were selected using the FindVariableFeatures function with mean greater than 0.0125 or less than 3 and dispersion greater than 0.5. These genes were used in performing the linear dimensionality reduction.

#### Clustering and Finding Markers

Principal component analysis was performed using the top 3000 most variable genes prior to clustering and number of the first principal components (PCs) were used based on the ElbowPlot as described below for different datasets. Clustering was performed using the FindClusters function which works on K-nearest neighbor (KNN) graph model with the granularity (resolution) ranging from 0.1-1.5. The datasets were projected as t-SNE plots.

### Statistical analysis

All numerical data analysis except for scRNA-seq data analysis was performed on Graphpad Prism8 software. Values were log-transformed to normalize distribution and variances where necessary. Immune cell numbers and cytokine measurement were analyzed by 2-way ANOVA with Bonferroni multiple comparisons test. Log-transformed viral titers were analyzed by repeated measures two-way ANOVA with Bonferroni multiple comparisons test. Inflammation scores were analyzed by repeated-measures two-way ANOVA or mixed-effects analysis with Geisser-Greenhouse correction and Bonferroni’s multiple comparisons test. The Geisser-Greenhouse correction was used for inflammation scores to correct any violations of sphericity and to provide a more restrictive, stringent calculation of p values. ROS MFI was measured by unpaired two-tailed Student’s t-test. qPCR results were analyzed by one-way ANOVA with Tukey’s multiple comparisons test. A p<0.05 was considered statistically significant. No experimental data points were excluded from statistical analysis, including potential outliers. Mouse and sample numbers per group and experimental repeat information is provided in the figure legends. All data points represent individual biological replicates, and the ‘n’ for each group refers to biological replicates. No power calculations were performed to determine sample size; rather sample sizes were determined based on historical data.

### DATA AVAILABILITY

The published article includes partial data from a single cell RNA-sequencing dataset generated during this study. These data are available at Gene Expression Omnibus (GEO) (accession code GSE161336). The key to access this data is sngheyymjxqbjsn.

---

**Figure 1 - Supplement 1.**
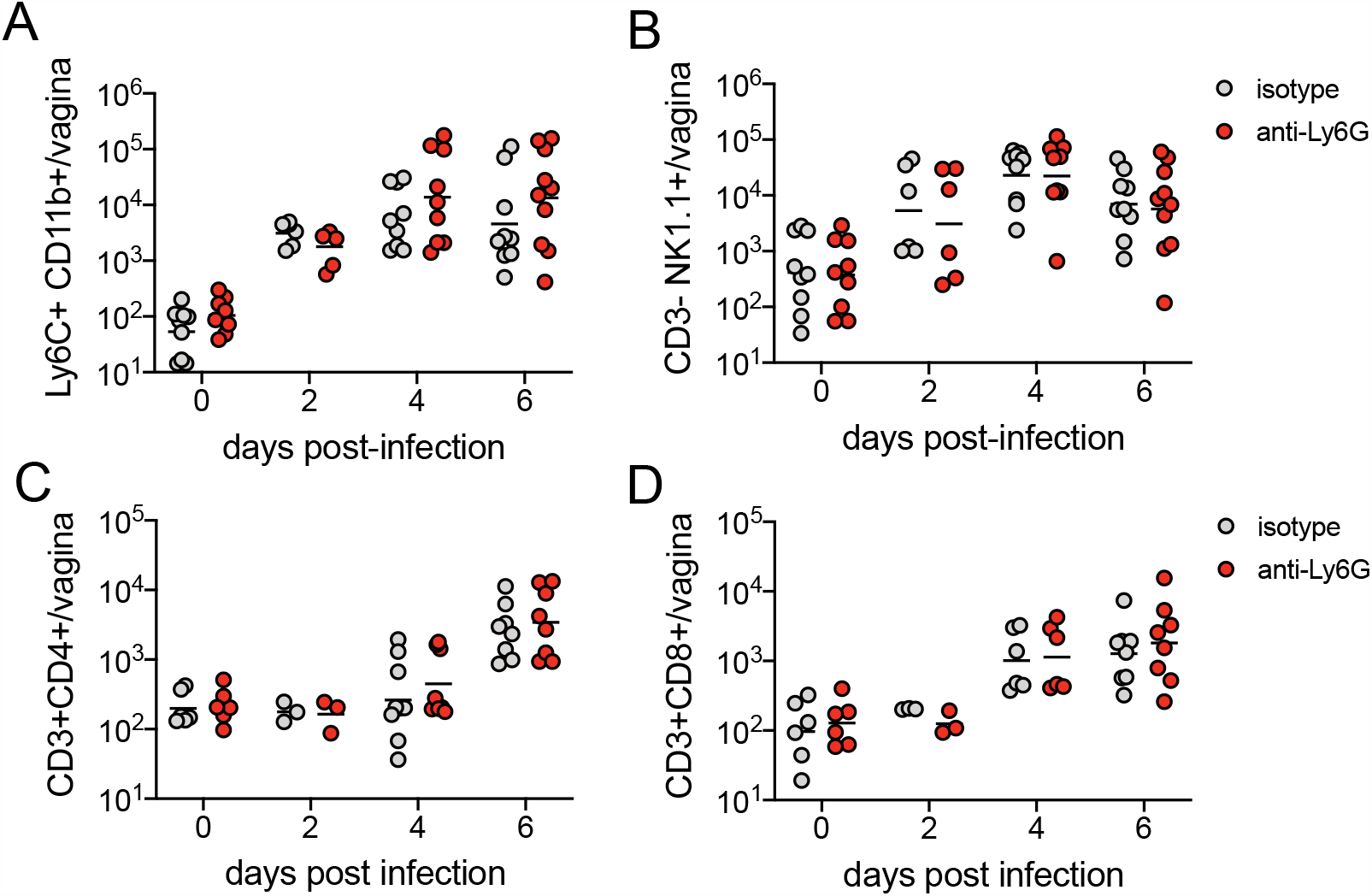
Neutrophil depletion does not affect magnitude of the immune cell response after HSV-2 infection. Mice were infected and treated as described in Figure 2. **A-D**. On the indicated d.p.i., immune cell infiltrates were measured in the vagina by flow cytometry. Ly6C+CD11b+ monocytes (**A**), CD3-NK1.1+ NK cells (**B**), CD3+CD4+ T cells (**C**) and CD3+CD8+ T cells (**D**) were enumerated in perfused tissue. (**A, B**) Isotype controls: n=6-9, anti-Ly6G: n=6-10. (**C, D**) Isotype controls: n=3-8, anti-Ly6G: n=3-9. All data are pooled from 2 independent experiments. Horizontal bars show mean (**A-D**). Statistical analysis was performed by two-way ANOVA with Bonferroni’s multiple comparisons test. All comparisons were not significant. Raw values for each biological replicate, epsilon values and specific p values are provided in Figure 1 - Supplement 1 source data.

**Figure 1 - Supplement 2.**
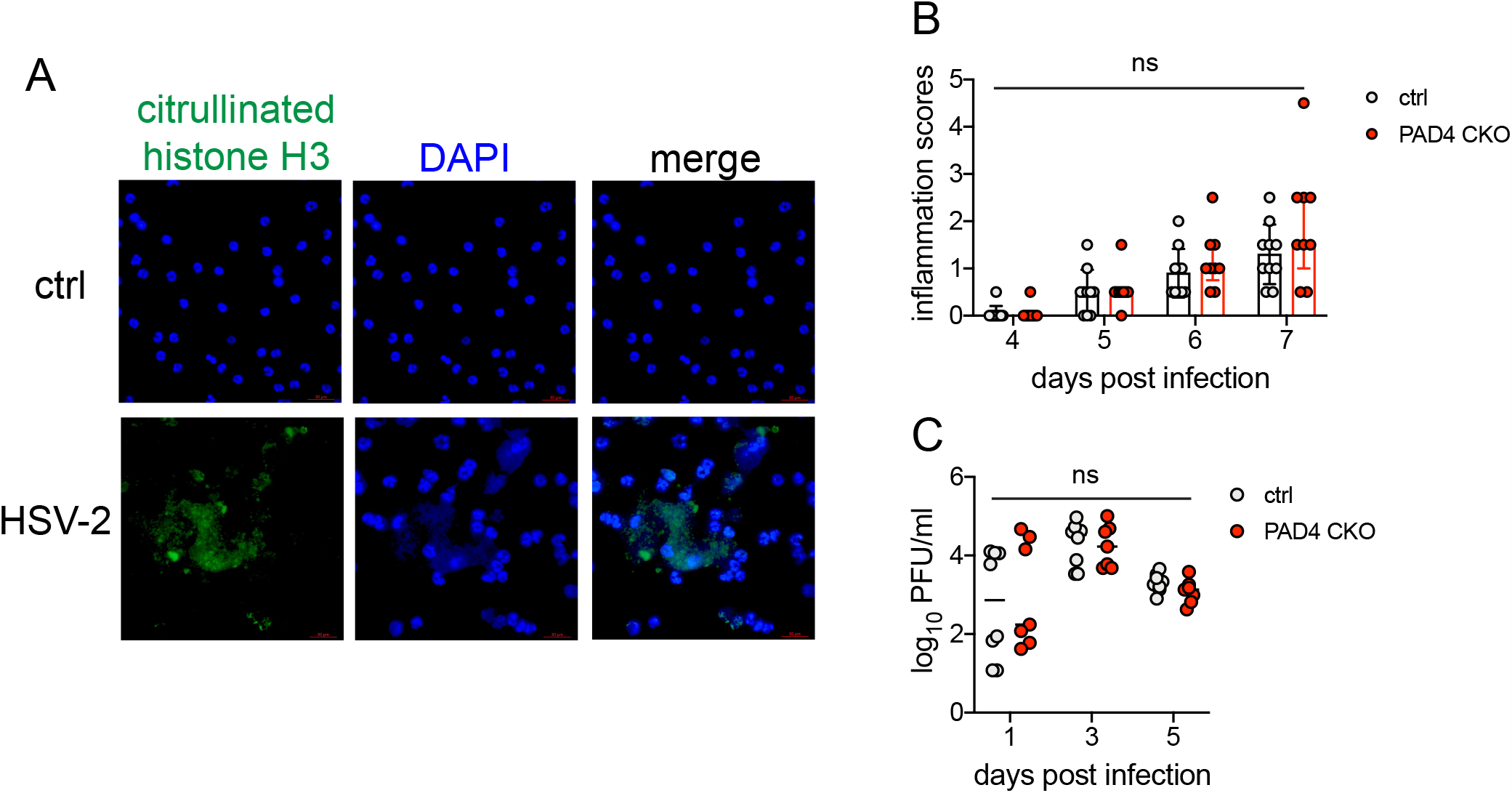
PAD4 is not required for development of genital inflammation during HSV-2 infection. **A.** Neutrophils were isolated from the bone marrow of naive C57BL/6 female mice and stimulated with heat-killed HSV-2 at an MOI of 1 for 4 hours (bottom row) or left unstimulated (top row). NETs are identified by areas of diffuse DAPI staining (blue) that overlap with citrullinated histone H3 (green). Data are representative of 2 independent experiments. **B-C**. Pad4^fl/fl^ x MRP8-Cre (PAD4 CKO) or Cre-littermate controls were infected with HSV-2 as described in Figure 1. Inflammation scores were monitored for 7 d.p.i. (Ctrl: n=10, KO: n=9) (**B**), and infectious virus was measured by plaque assay in vaginal washes collected on the indicated d.p.i. (Ctrl: n=8, CKO: n=8) (**C**). Data in **B-C** is pooled from 2 independent experiments. Statistical significance was analyzed by repeated measures two-way ANOVA with (**B**) or without (**C**) Geisser-Greenhouse correction and Bonferroni’s multiple comparisons test, ns = not significant. Raw values for each biological replicate, epsilon values and specific p values are provided in Figure 1 - Supplement 2 source data.

**Figure 1 - Supplement 3.**
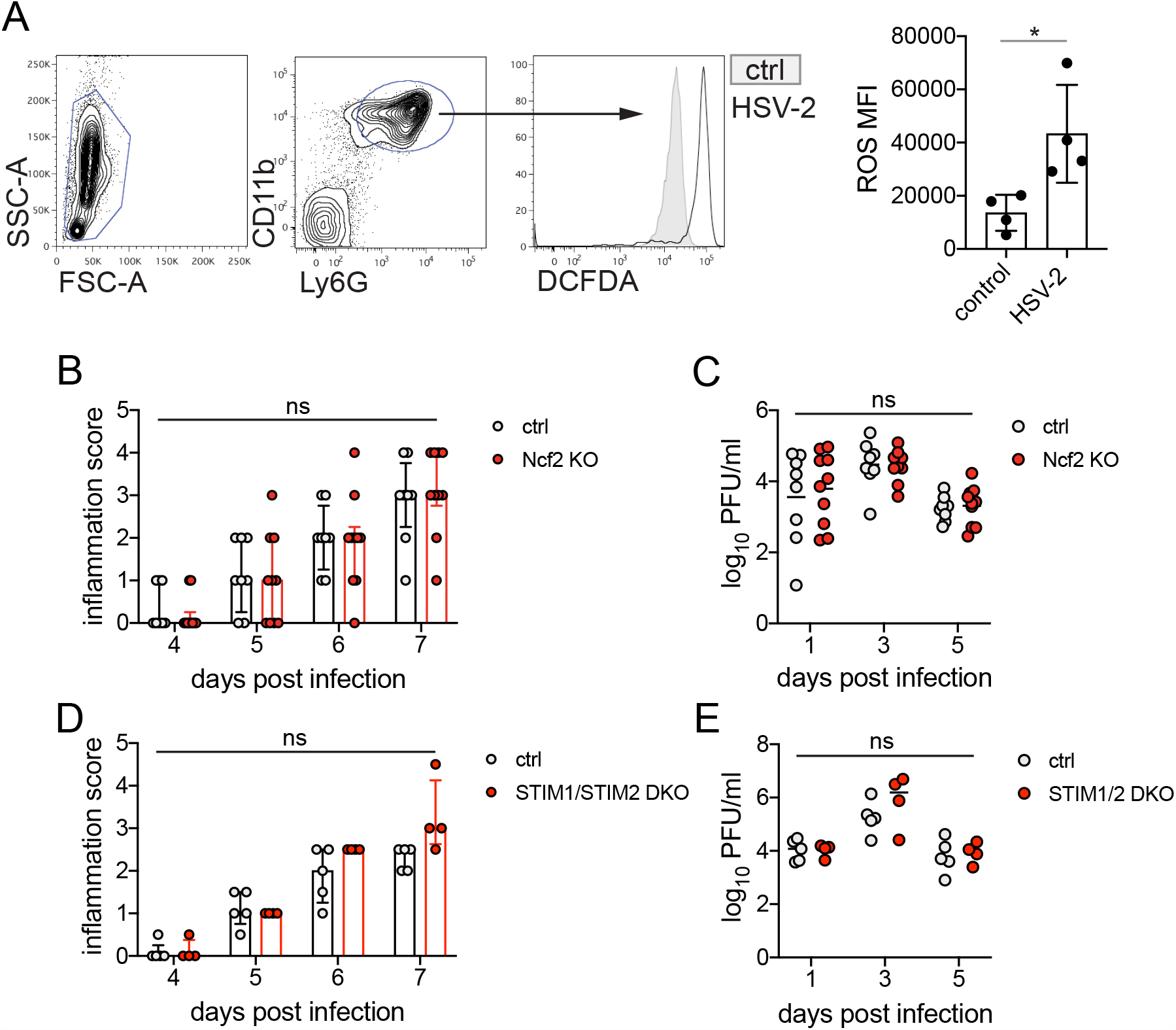
ROS production and STIM1/STIM2 expression in neutrophils are not required for genital inflammation after HSV-2 infection. **A.** Neutrophils were isolated from the bone marrow as using a Histopaque gradient and stimulated with heat-killed HSV-2 at an MOI of 5 for 16 hours (n=4) of left unstimulated (n=4) and then incubated with DCFDA. Plots show gating for leukocytes (left) and CD11b+ Ly6G+ neutrophils (middle). DCFDA fluorescence was measured by flow cytometry in unstimulated (shaded histogram) or HSV-2 stimulated (open histogram) neutrophils. Graph shows mean fluorescence intensity (MFI) of DCFDA (ROS). Ncf2 deficient mice (n=10) or littermate controls (n=8) (**B-C**) or STIM1/STIM2 deficient mice (STIM1/STIM2 DKO, n=4) or littermate controls (n=5) (**D-E**) were infected as described in Figure 1. Mice were monitored for genital inflammation for one week after infection **(B, D)**. Infectious virus was measured by plaque assay in vaginal washes collected on the indicated days (**C, E**). Bars in **A** show mean with SD. Bars in **B** and **D** show median with interquartile range. Bars in **C** and **E** show mean. Data in **A-C** are pooled from 2 independent experiments, data in **D-E** was performed once. Statistical significance was analyzed by unpaired Student’s t-test (**A**), repeated measured two-way ANOVA with (**B, D**) or without (**C, E**) Geisser-Greenhouse correction and Bonferroni’s multiple comparisons test, *p<0.05, ns = not significant. Raw values for each biological replicate, epsilon values and specific p values are provided in Figure 1 - Supplement 3 source data.

**Figure 2 - Supplement 1.**
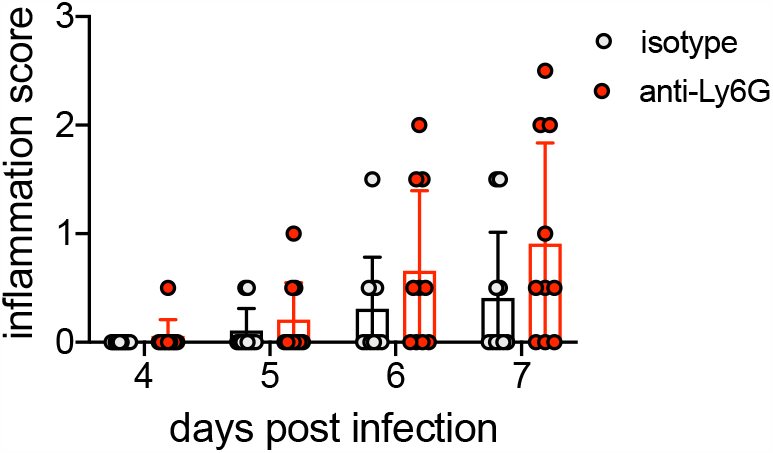
Neutrophil depletion prior to HSV-1 genital infection has little impact on disease progression. C57BL/6J females were infected with 10^4^ PFU HSV-1 McKrae and treated with neutrophil-depleting antibodies as described in Figure 1. Disease progression was measured during the first 7 d.p.i. for anti-Ly6G (n=10) and isotype control (n=10) mice. Bars show median and interquartile range. Data are pooled from 2 independent experiments. Statistical significance was measured by repeated measured two-way ANOVA with Geisser-Greenhouse correction and Bonferroni’s multiple comparisons test. All comparisons were not significant. Raw values for each biological replicate, epsilon values and specific p values are provided in Figure 2 - Supplement 1 source data.

**Figure 3 - Supplement 1.**
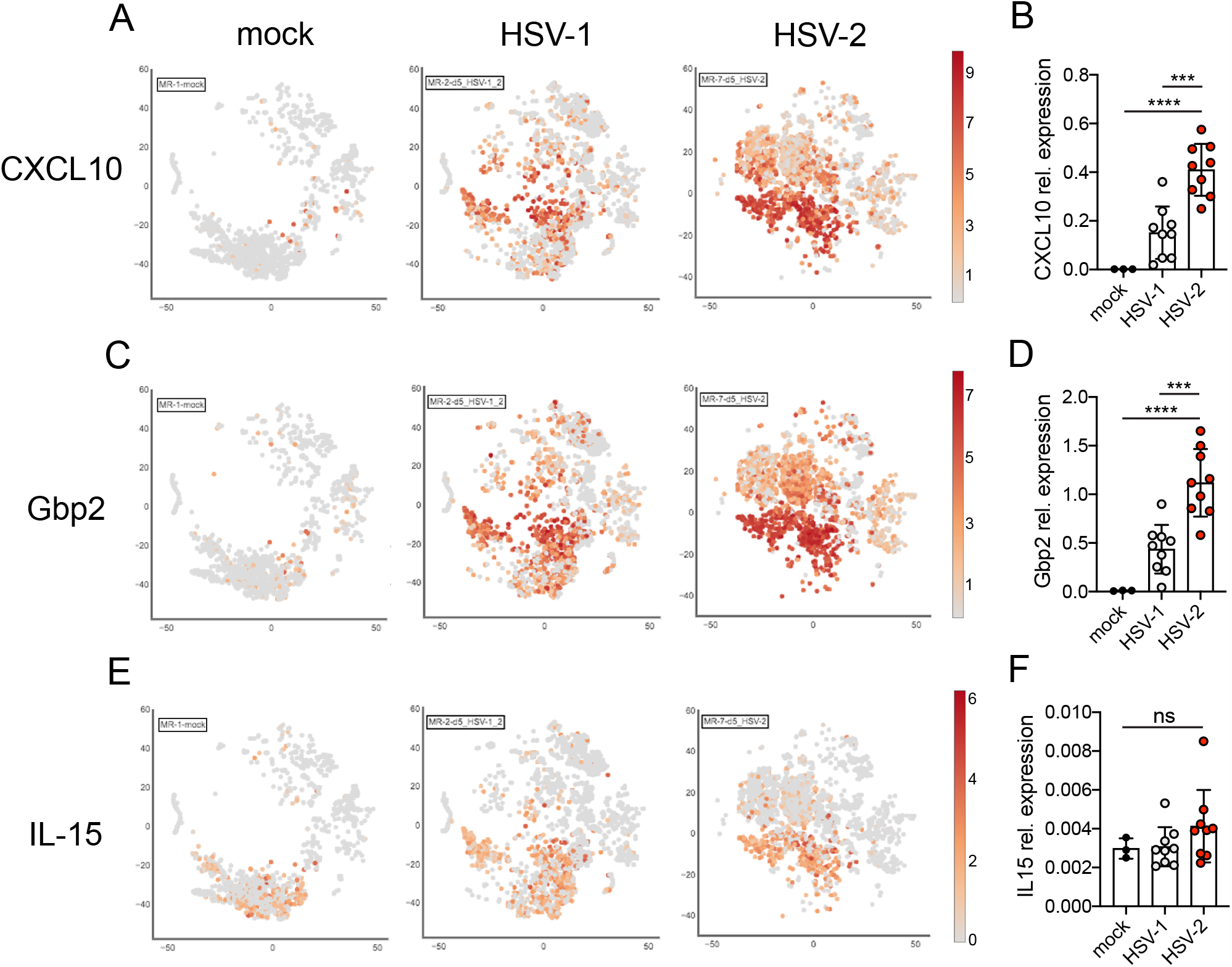
Validation of ISG expression in the vagina. Expression of CXCL10 (A, B), Gbp2 (C, D) or IL-15 (E, F) in the vagina of mock inoculated mice or mice at 5 d.p.i. with HSV-1 or HSV-2. tSNE visualization of select ISG transcripts in live vaginal cells profiled by single cell RNA-seq (A, C, E). Expression of the same ISGs relative to Rpl13 was measured by qRT-PCR in whole vaginal tissue harvested from mock inoculated mice (n=3) or mice at 5 d.p.i. with HSV-1 (n=9) or HSV-2 (n=9) (B, D, F). Bars in B, D and F show mean and SD. Data are pooled from 2 independent experiments. Statistical significance was measured by one-way ANOVA with Tukey’s multiple comparisons test. ***p<0.005, ****p<0.001, ns = not significant. Raw values for each biological replicate and specific p values are provided in Figure 3 - Supplement 1 source data.

**Figure 3 - Supplement 2.**
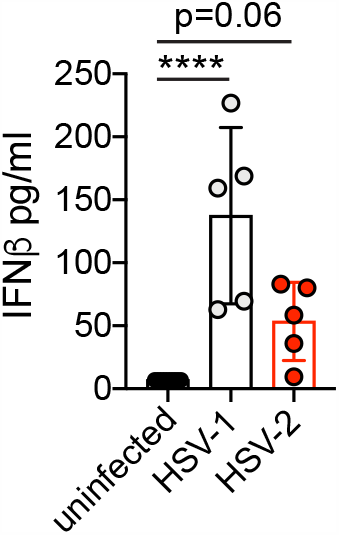
Type I IFN is robustly produced in the vagina early after acute HSV-1 or HSV-2 infection. C57BL/6J mice were infected as described in Figure 2. Type I IFN was measured by ELISA in vaginal washes collected from uninfected mice (n=10), or mice at 2 d.p.i. with HSV-1 (n=5) or HSV-2 (n=5). Bars show mean with SD. Experiment was performed once. Statistical significance was measured by one-way ANOVA with Dunnett’s multiple comparisons test. ****p<0.001. Raw values for each biological replicate, epsilon values and specific p values are provided in Figure 3 - Supplement 2 source data.

**Figure 4 - Supplement 1.**
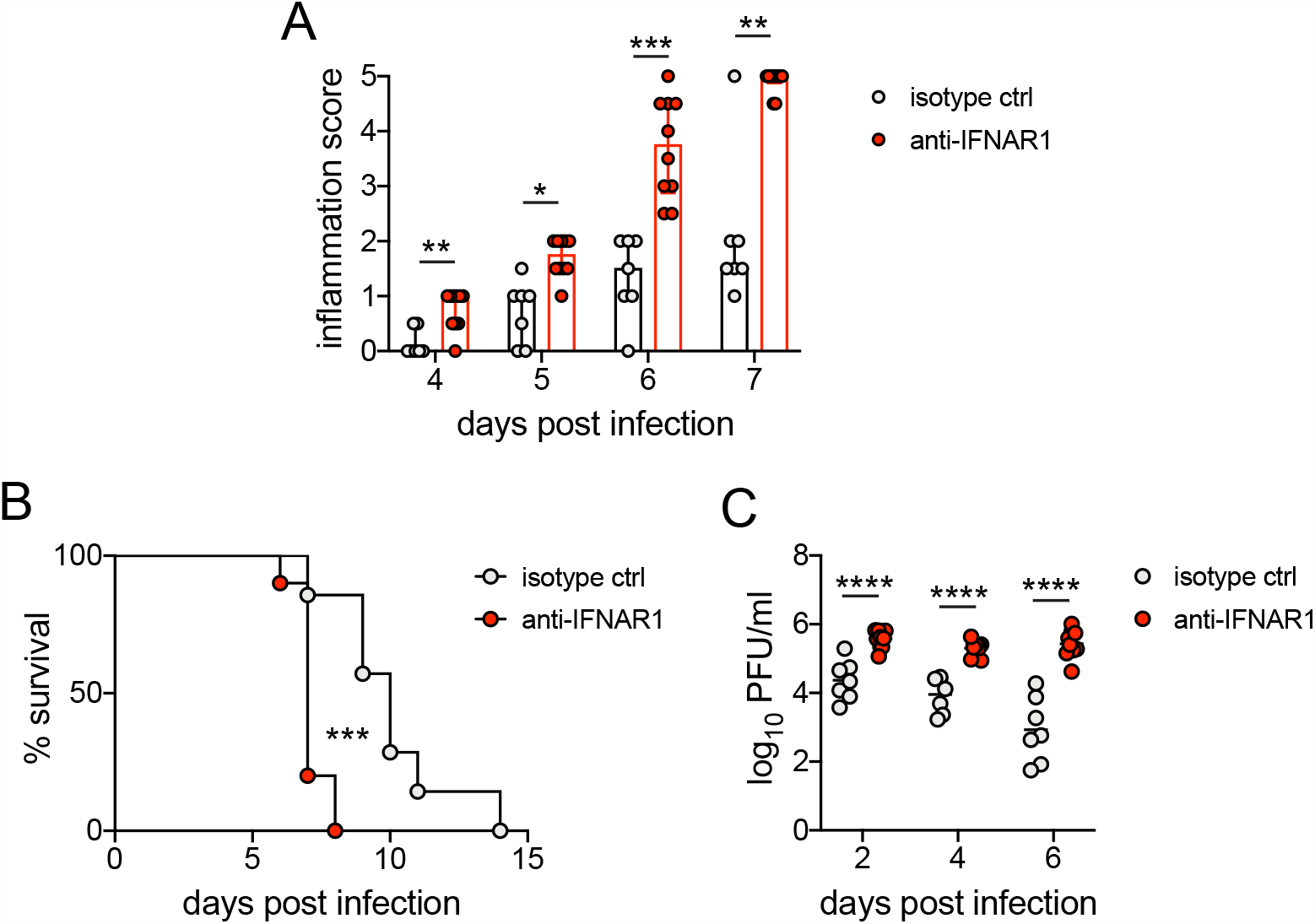
Early blockade of IFNAR1 leads to accelerated and more severe disease after HSV-2 infection. C57BL/6J mice were infected as described in Figure 1. On the day of inoculation, mice were injected i.p. with 1mg anti-IFNAR1 antibody (n=10) or an isotype control (n=7). **A**. Inflammation scores of anti-IFNAR antibody or isotype control treated mice for the first 7 d.p.i.. **B**. Survival of mice over the course of two weeks. **C**. Infectious virus as measured by plaque assay in vaginal washes collected at the indicated d.p.i.. Bars in **A** show median and interquartile range, bars in **C** show mean. Data are pooled from 3 independent experiments. Statistical significance was measured by repeated measured two-way ANOVA with Geisser-Greenhouse correction and Bonferroni’s multiple comparisons test (**A**), log-rank test (**B**) and mixed-effects analysis with Bonferroni’s multiple comparisons test (**C**). *p<0.05, **p<0.01, ***p<0.005, ****p<0.001. Raw values for each biological replicate, epsilon values and specific p values are provided in Figure 4 - Supplement 1 source data.

**Figure 6 - Supplement 1.**
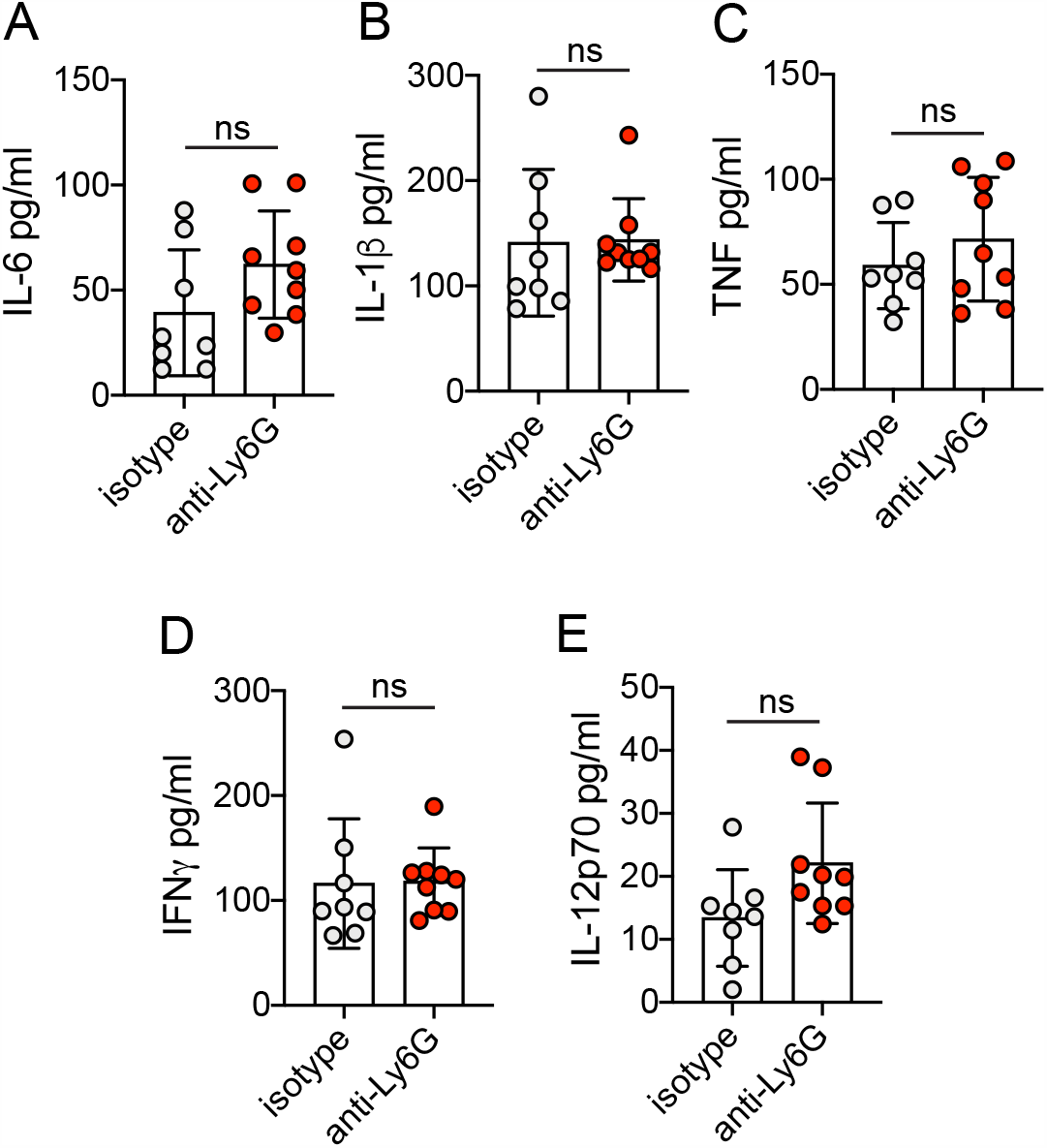
Neutrophils do not control production of common pro-inflammatory and antiviral cytokines during HSV-2 infection. Vaginal washes were collected at 5 d.p.i. from mice that were infected and treated as described in Figure 1. IL-6 (A), IL-1b (B), TNF (C), IFNγ (D) and IL-12p70 (E) were measured by multiplexed Bioplex assay. Isotype controls: n=8, anti-Ly6G: n=9. All data are pooled from 2 independent experiments. Statistical analysis was performed by unpaired t-test. ns = not significant. Raw values for each biological replicate and specific p values are provided in Figure 6 - Supplement 1 source data.

